# Unidirectional Single-File Transport of Full-Length Proteins Through a Nanopore

**DOI:** 10.1101/2021.09.28.462155

**Authors:** Luning Yu, Xinqi Kang, Fanjun Li, Behzad Mehrafrooz, Amr Makhamreh, Ali Fallahi, Joshua C. Foster, Aleksei Aksimentiev, Min Chen, Meni Wanunu

## Abstract

Nanopore sensors could revolutionize single-molecule proteomics by providing a means for identification of known proteins through fingerprinting or by de novo sequencing. However, the complex chemical and physical properties of proteins present multiple challenges to the conventional nanopore sensing method, predominantly, the single-file threading of a protein chain into a nanopore and its transport through it. Herein we describe a general approach for realizing unidirectional transport of full-length proteins through nanopores. We show that the combination of a chemically resistant biological nanopore platform and a high concentration guanidinium chloride buffer enables protein unfolding and unidirectional transport through a pore, propelled by an electroosmotic effect that largely owes to the guanidinium chloride presence. The uniform and slow (~10 µs/amino acid) single-file transport, when combined with supervised machine learning of the electrical current signatures obtained, allows us to use to discern the protein threading orientation and identity. In conjunction with a method for tail-modification of native proteins and higher-resolution nanopores, our approach could offer a path towards direct single-molecule protein fingerprinting without the requirement of a motor enzyme.

## Introduction

Living systems use complex molecules to encode and carry out their essential functions, with nucleic acids being the central information carrier across generations and within species. Tremendous technological progress has been made in recent years in high-throughput and long-read genomic sequencing^1^, including several single-molecule techniques^2^, where the nucleotide sequence of individual DNA molecules is determined by either monitoring DNA replication in real-time^3^ or by passing a DNA strand through a nanopore detector^4,5^. While DNA serves as a blueprint for the type of biomolecules that a cell can produce, genomic information alone is not sufficient to predict a cell’s phenotype^6,7^, which is ultimately defined by the type and abundance of cellular proteins.

More than 10^4^ protein types make up the canonical human exome, with a multitude of protein isoforms^8,9^ and post-translational modifications (PTMs)^10,11^ giving a cell’s phenotype its functionality. Molecular characterization of these diverse phenotypes requires quantitative methods for protein counting, sequencing, and discrimination among various isoforms and PTMs^12^. Mass spectrometry (MS) is currently the gold standard method for these characterizations, which commonly require protein fragmentation, quantification, and sequence reconstruction. The high sensitivity of MS to minute protein quantities permits single-cell proteomics^13^. However, low peptide ionization rates and other limitations result in only a few percent sampling efficiencies^14,15^. Alternative approaches are needed to deliver a complete single-cell proteome without extensive fragmentation.

Nanopores have emerged as key components of new proteomics tools^16^. The methodology of nanopore DNA sequencing has been adapted to sense the amino acid composition of model peptides and proteins^16–21^. However, biological proteins are much more complex than DNA, given their diverse secondary structures, strong intramolecular interactions, and a heterogeneous distribution of the electrical charge along their polypeptide chains. Voltage^22–25^, temperature^26^, and chemical denaturation ^18,27–32^ have been used to enhance the access of unfolded proteins to the nanopore constrictions. However, protein unfolding does not guarantee protein translocation because an unmodified proteins cannot be electrophoretically driven through a nanopore in the same manner as a charged DNA strand can^33^. Enzyme-assisted unfolding and translocation of large proteins have been demonstrated using ClpXP as a motor to linearize and pull the protein through α-hemolysin^34,35^. Recent works using phi29 DNA polymerase^36^ or Hel308 DNA helicase^37,38^ have shown enzyme-mediated ratcheting or unwinding of peptide-DNA conjugates through the MspA nanopore. However, to date, enzyme-free readout of a full-length protein using a nanopore has not been achieved.

Here, we demonstrate an enzyme-free platform for single-file unidirectional transport of full-length proteins through a nanopore reader. Electroosmotic flow, enhanced by the use of GdmCl as a denaturant, drives protein transport through the nanopore, conferring uniform and slow (~10 µs/amino acid) single-file protein transport. The ionic current signals produced by the transport are found to carry the information about the protein sequencing, which we demonstrate by matching the signals produced by the N-terminus and C-terminus transport of the same protein, the transport of a double concatemer of the same protein, and by determining the composition of a binary protein mixture. With further development, our approach paves the way for single-molecule protein identification and quantification of the protein isoforms.

## Results and Discussion

### Experimental setup for unfolding and transporting full-length protein through a nanopore reader

Our experimental setup (**Figure 1A**) comprises a wedge-on-pillar (WOP) membrane support^39^, a poly(1,2-butadiene)-b-poly(ethylene oxide) (PBD_n_-PEO_m_) block-copolymer bilayer that spans the aperture, as well as an inserted wild-type α-hemolysin channel, which is the nanopore used in our experiments. We chose this block copolymer membrane for its chemical compatibility with GdmCl buffers, and high voltage tolerance (>350 mV for 100 μm diameter bilayer membranes) when combined with the WOP support^40^. Also depicted in the figure is our use of high concentration GdmCl buffer, critical for protein unfolding. **Figure 1B** shows current vs. voltage curves for single α-hemolysin channels at different buffer conditions. All curves exhibit significant asymmetry, with higher current amplitudes at positive voltages. The impact of GdmCl on noise in α-hemolysin is moderate: The 10 kHz bandwidth noise at 300 mV is 7.2 pA for 2.5 M KCl and 10.7 pA for 1 M KCl + 2.0 M GdmCl, respectively. Power spectra at 0 mV and 300 mV for these two buffers (see inset) indicate a slight increase in the noise in the low-to-intermediate frequency regime (<5 kHz).

**Figure 1.**
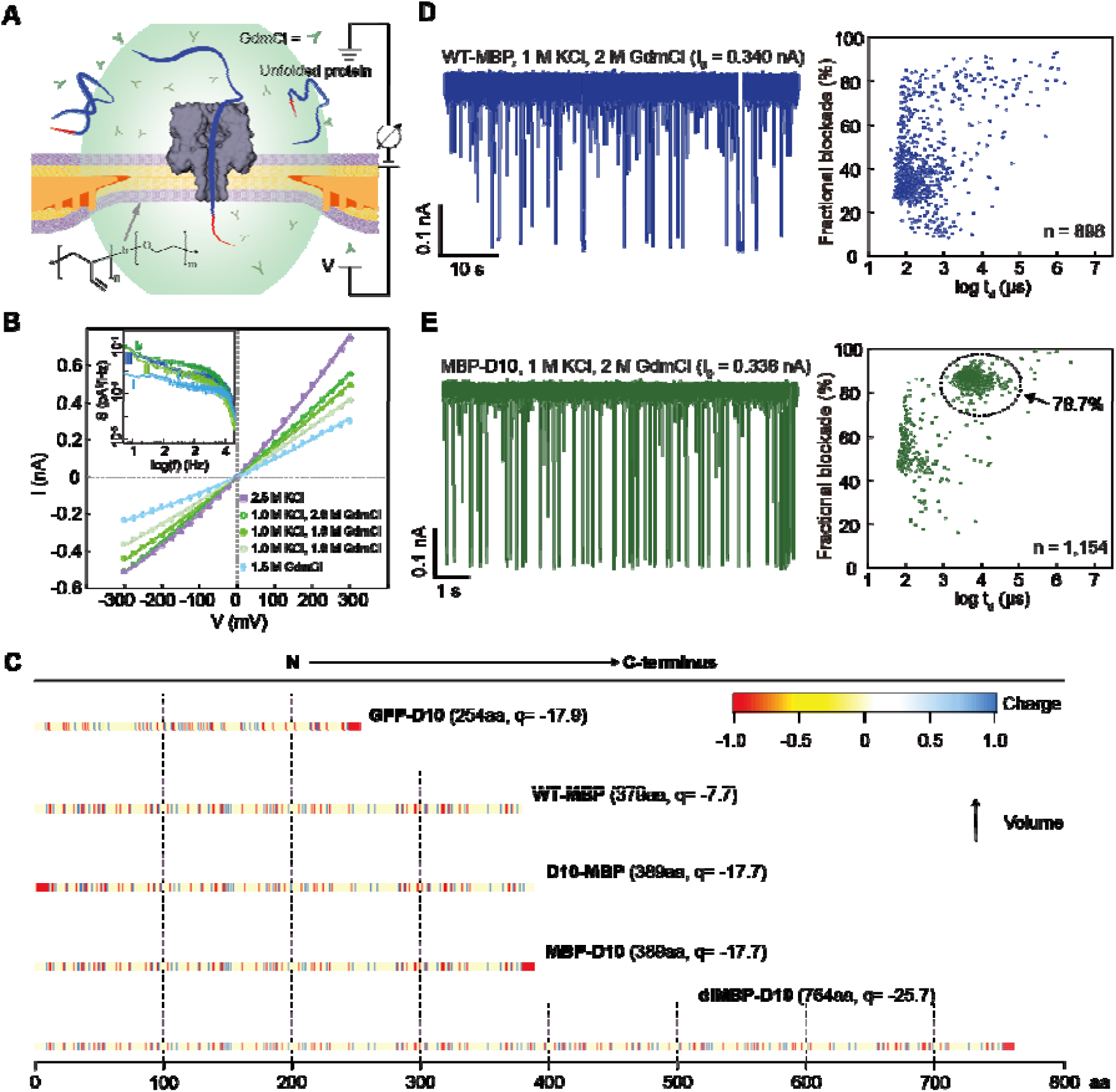
Enzyme-free full-length protein translocation through nanopores. **A)** Schematic cut-away view of a PBD_n_-PEO_m_ block-copolymer bilayer (inset shows polymer structure) suspended on the WOP aperture (orange depicts lipid solvent), with an α-hemolysin nanopore inserted into the bilayer. The guanidinium chloride (GdmCl) buffer unfolds the analyte proteins while leaving the α-hemolysin nanopore intact. A pair of Ag/AgCl electrodes generate transmembrane voltage and measure the ionic current through the nanopore. **B)** Current-voltage dependence of a single α-hemolysin channel at several buffer conditions (*V* is applied to the *trans* chamber). **C)** Graphical representations of the formal charges at pH 7.5 for the protein constructs used in this study in their unfolded (solvent accessible) state. A plot above each charge graph shows the relative amino acid volumes (pink). **D)** Current vs. time trace (left) and the fractional blockade vs. dwell time scatter plot (right) recorded from wild-type maltose binding protein (WT-MBP) at *V* = 175 mV, [WT-MBP] = 0.35 µM. Open pore current (I_o_) is indicated above each trace. **E)** Same as in panel D but for MBP containing a C-terminus aspartate tail (MBP-D10), [MBP-D10] = 0.35 µM. The scatter plot indicates the fraction of detectable events near the dashed circle.

Using the above experimental setup, we conducted nanopore translocation experiments on variants of the α-helix-rich maltose-binding protein (MBP) in its monomeric form (denoted as either MBP-D10 or D10-MBP, depending on the C- or N-terminus attachment of the D_10_ tail) and its dimeric form (diMBP-D10, with a GGSG linker between two MBP monomers). The nanopore translocation experiments were repeated using green-fluorescent protein (GFP), which stable β-barrel structure makes this protein notoriously challenging to unfold. **Figure 1C** shows the net charge (at pH 7.5)^41^ and length (number of aa’s) of each protein, as well as pKa-based graphical profiles of the charge^41^ and relative volume^42^ of each aa residue.

The impact of a charged tail on protein threading is indicated in **Figure 1D** and **1E**, where current traces for wild-type MBP (WT-MBP) and MBP-D10 are shown. While in both experiments, protein concentration was 350 nM, the capture rates for MBP-D10 (9.3 s^−1^µM^−1^) were ~80% higher than for WT-MBP (5.2 s^−1^µM^−1^). Further, WT-MBP events had a broad range of amplitudes and short durations (<1 ms), whereas MBP-D10 had nearly 79% of the events forming a tight distribution characterized by a ~85% fractional blockade and dwell times between 1 to 10 ms. Such reproducible dwell times and current amplitudes suggest an efficient and deterministic translocation process mediated by the insertion of the D_10_ tail, which is in agreement with earlier reports where charged tags, such as DNA oligos^24^ or poly-aspartate tails^34^, were used to direct protein capture. However, modifying a full-length protein without any prior genetic engineering is crucial for a path to analysis of native proteins. We therefore demonstrate that DNA conjugation to a full-length protein tail is possible, albeit at low yields (~5-6%), by labeling WT-MBP with a dT_20_ DNA oligo (see **SI, Figure S1** and **Supplementary Note 1**). We confirmed that the addition of the dT_20_ tail enhances threading of MBP through the pore, as indicated by an enriched cluster of events with similar dwell times and current blockades as observed for MBP-D10 (compare **Figure 1E** to **SI Figure S2**).

## Influence of GdmCl concentration on complete protein unfolding

Bulk measurements suggest that MBP has an unfolding midpoint at 1.0 M GdmCl at room temperature, and that MBP fully denatures when GdmCl concentration exceeds ~1.2 M^43,44^. After adding MBP-D10 to the *cis* chamber, in **Figure 2A**, the current blockade vs. dwell time scatter plots (for 1.0 M, 1.5 M, and 2.0 M GdmCl, respectively) show two main distributions, highlighted by dashed red and blue circles, in addition to a very “fast” population at ~100 µs, which most likely corresponds to protein collisions with the pore entrance. Consistent with the scatter plots, the current traces in **SI Figure S4** show two types of events (long and short). We assign the long-lived events (red dashed circle) to population P_F_, which encompass events from partially folded proteins. For the reasons described in the next paragraph, we ascribe the tight, shorter-lived population P_L_ (blue dashed circle) to events produced by linearized, completely unfolded proteins. Since these two populations are generally well-resolved, we have determined the percentage of linear protein events P_L_ as a function of GdmCl concentration (see **SI, Figure S17**). As shown in **Figure 2A**, P_L_ increases with the GdmCl concentration from ~36% at 1.0 M GdmCl to >93% at 2.0 M GdmCl. The existence of one population at 2.0 M GdmCl suggests that the protein is fully unfolded during its translocation through the pore.

**Figure 2.**
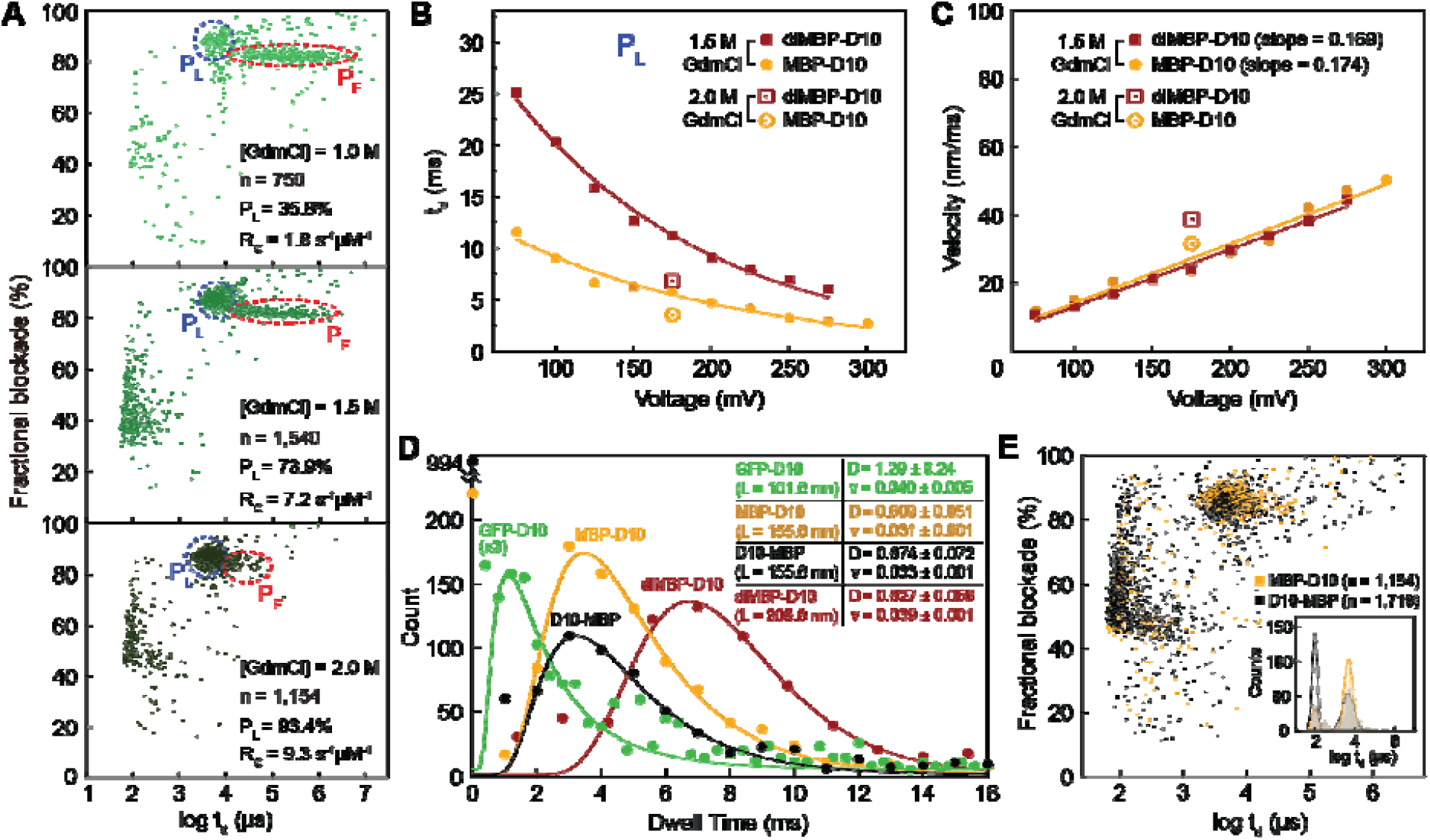
Transport properties of unfolded protein analytes. **A)** Fractional current blockade vs. dwell time scatter plots for MBP-D10 in 1.0 M, 1.5 M, and 2.0 M GdmCl buffer (+1 M KCl). Red and blue ovals show populations that correspond to P_F_ (partially folded) and P_L_ (linear or unfolded) states of MBP-D10, respectively. [MBP-D10] = 0.7 µM for 1.0 M GdmCl and 0.35 µM for 1.5, 2.0 M GdmCl experiments. **B)** Mean dwell time vs. voltage with exponential fitting for the P_L_ populations of MBP-D10 and diMBP-D10, respectively (error bars represent the FWHM of the distribution fits). **C)** Protein transport velocities calculated from estimated protein contour length and observed dwell times as a function of applied voltage (error bars are based on the dwell time distribution widths shown in **B**). Buffer conditions for data shown in **B and C** are 10 mM Tris, pH 7.5, 1 M KCl, and either 1.5 or 2.0 M GdmCl. For the latter, data at only one voltage (175 mV) are shown. **D)** Dwell time histograms for GFP-D10, MBP-D10, D10-MBP, and diMBP-D10 along with the mean diffusion coefficients (nm^2^/µs) and velocities (nm/µs) determined from fits to the 1D Fokker-Planck equation^55,79,80^. **E)** Fractional current blockade vs. dwell time scatter plots and dwell time histograms for C-terminus (MBP-D10) vs. N-terminus (D10-MBP) threading and transport of full-length MBP. Experiments in **D** and **E** were performed in 1 M KCl, 2.0 M GdmCl, 10 mM Tris, pH 7.5, under a 175 mV bias applied to the *trans* chamber.

## Evidence of steady voltage-driven protein translocations

**Figure 2B** plots the mean dwell time of the P_L_ population as a function of voltage for experiments conducted using an MBP monomer (MBP-D10) or a dimer (diMBP-D10) (see **SI Figure S11** for the dwell time histograms). The average dwell time of population P_L_ decreases as the voltage increases. Importantly, the dwell time for the MBP dimer (di-MBP-D10) is a factor of two larger than that for the MBP monomer (MBP-D10). In contrast, the dwell time of events attributed to partially folded proteins, P_F_, are much larger for both MBP-D10 and diMBP-10 than for the unfolded events at all voltages and exhibit a steeper voltage dependence than the P_L_ population (**SI Figure S12**). The plot of the protein mean “velocity”, **Figure 2C**, calculated by dividing the protein contour length (0.34 nm per amino acid) by the dwell time (from **Figure 2B**) shows a linear dependence on voltage for both monomeric and dimeric MBP, thus indicating that the velocity does not depend on protein length. The electrophoretic mobility of a protein is described by

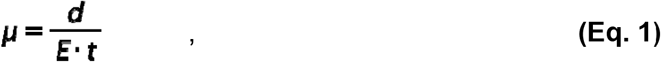

where *d* is the protein’s contour length, and d / t = ***v***_e_ is the protein translocation velocity^45^. Estimating the electric field ***E*** as the ratio of the voltage to the length of the α-hemolysin lumen, ***D*** = 5 nm, we obtain

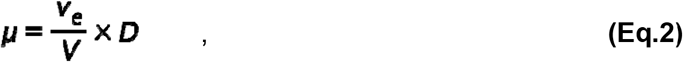

where ***v***_***e***_ **/*V*** is the slope of the fits from **Figure 2C**. Using Eq. 2, we find the electrophoretic mobility of MBP-D10 and diMBP-D10 at μ_m_ = 8.70 × 10^−9^ ***cm***^**2**^ **/ *V* ·*s*** and μ_d_ = 8.45 × 10^−9^ ***cm***^**2**^ **/ *V* ·*s***, respectively. Moreover, two extra datapoints at 175 mV are shown for 2 M GdmCl condition (**Figure 2C**), indicating faster translocating speeds. The uniform translocation speed, and its linear relationship with voltage resembles ssDNA translocation through α-hemolysin^46–48^. Finally, a plot of the event rates (See **SI, Figure S18**) reveals a low-voltage regime characterized by short-lived high-frequency collisions and an exponentially increasing capture rate at higher voltages (V > 150 mV), which suggests an entropic barrier for capture^49^.

In **Figure 2D**, we present dwell-time distributions for GFP-D10 (254 aa), MBP-D10 (389 aa), D10-MBP (389 aa), and diMBP-D10 (764 aa) obtained at 175 mV applied voltage and 2 M GdmCl denaturant concentrations. We found no dependence of the protein transport time on its orientation of entry (MBP-D10 vs. D10-MBP), which differs from the orientation dependence of DNA transport through α-hemolysin^50^. This is supported by **Figure 2E**, which shows current blockade vs. dwell time scatters for MBP-D10 and D10-MBP. To a large extent, there is an overlap in the dwell time distributions, with the exception of D10-MBP that is captured by the pore less effectively, as indicated by many collisions of shorter dwell times (~100 µs) and lower current blockades. However, it is noteworthy that protein translocation from either direction proceeds with the same speed. Drift velocities for all molecules in this study are in the range of 0.031 – 0.04 nm/µs. The extended backbone distance between amino acids in a protein chain (0.34 nm)^51^ translates to a mean residence time of ~10 µs per amino acid in the pore. This average translocation velocity is roughly an order of magnitude smaller than that measured for a ssDNA transport through α-hemolysin (0.15 nm/µs) ^46^.

## Electroosmotic flow drives protein transport

To determine how Gdm^+^ ions enable unidirectional transport of unfolded peptides, we built seven all-atom systems, each containing a different 52-residue fragment of the MBP protein (Table S1) threaded through α-hemolysin, a lipid membrane, and 1.5 M GdmCl/1 M KCl electrolyte (**Figure 3A**). For comparison, two variants of each system were built, differing by the electrolyte solution composition (1.5 M GdmCl and 2.5 M KCl). Each design was equilibrated using the all-atom MD method^52^ and then simulated under a +200 mV bias for approximately 1,500 ns (see Materials and Methods for details).

**Figure 3.**
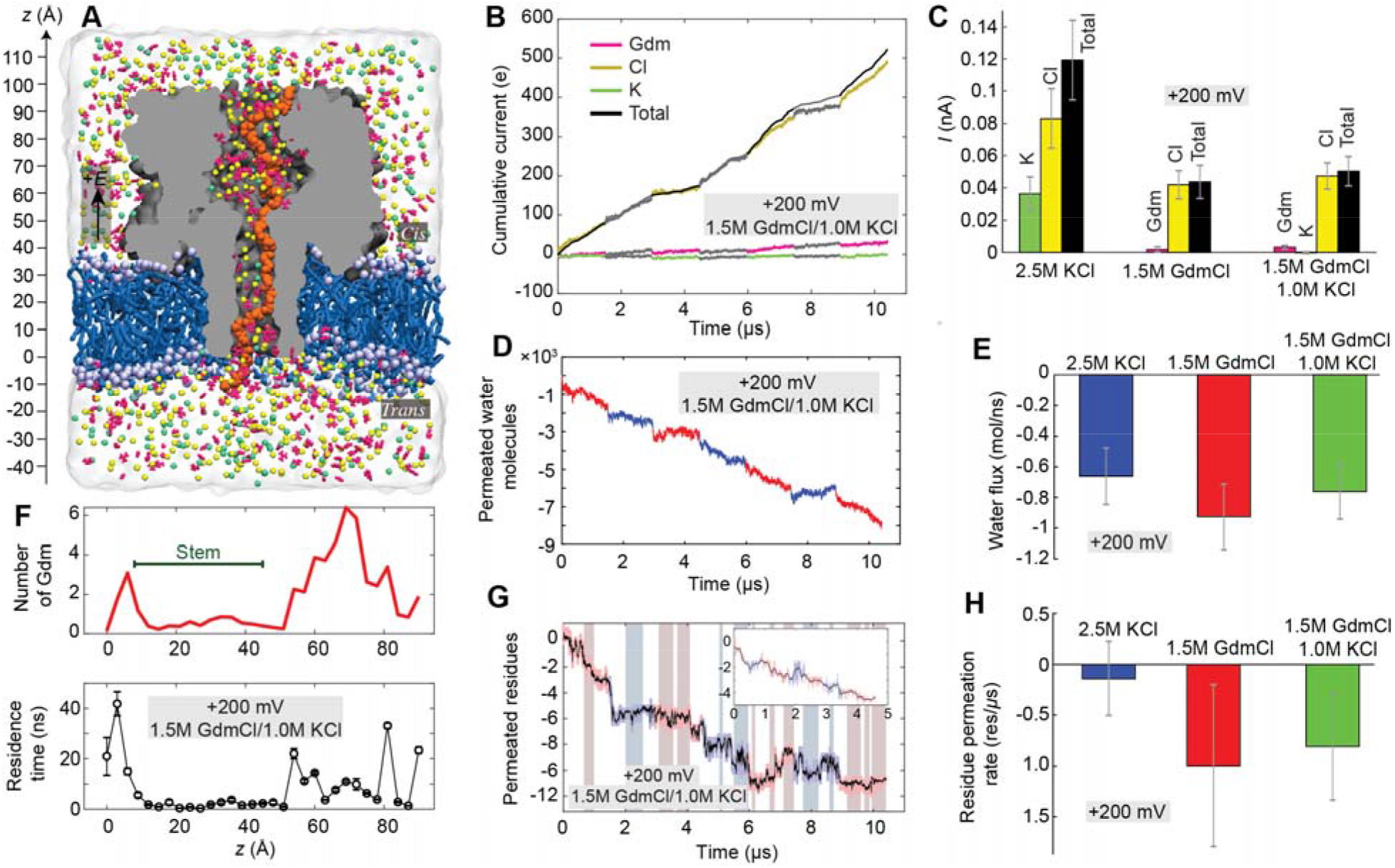
MD simulation of ion, water, and peptide transport through α-hemolysin. **A)** All-atom model of α-hemolysin (gray) containing a fragment of the MBP protein (orange), embedded in a lipid membrane (blue) and submerged in the 1.5 M GdmCl, 1 M KCl electrolyte mixture. **B)** Total charge carried by ion species in seven independent MD simulations differing by the sequence and initial conformation of the MBP fragment. Hereafter each trace is shown using two alternating colors to indicate data from independent trajectories. The traces are added consecutively to appear as a continuous permeation trace. The slope indicates the average current. **C)** Average ionic current for the three electrolyte conditions. Hereafter the average and the standard error are calculated considering each trajectory-averaged value as a result of an independent measurement. **D)** Number of water molecules permeated through the α-hemolysin constriction (residues 111, 113, and 147). Negative values indicate transport in the negative *z*-axis direction (defined in panel A). **E)** Average water flux for each electrolyte condition. **F)** The number of Gdm^+^ ions within 3 Å of the nanopore inner surface (top) and their average residence time (bottom) along the transmembrane pore of α-hemolysin. **G)** Number of amino acids permeated through α-hemolysin constriction under a +200 mV bias in the 1.5 M GdmCl/1.0 M KCl electrolyte simulations. Highlights indicate the parts of the trajectories where the peptide density within the stem (8<z<45 Å) of α-hemolysin is constant; the inset shows consecutive addition of the highlighted regions. **H)** the Average number of translocated amino acids for each electrolyte condition.

In the case of the GdmCl/KCl electrolyte, 94% of the blockade current was carried by Cl^−^ ions, whereas the current carried by Gdm^+^ and K^+^ ions was 6 and 0%, respectively (**Figure 3B**). Similarly, strong ionic selectivity was observed for pure GdmCl electrolyte (**Figure 3C**, and **SI Figure S20**). The ionic selectivity was less pronounced but still substantial (70 and 30% for Cl^−^ and K^+^ currents, respectively) for pure KCl. Consistent with the ion selectivity, we observed strong electroosmotic effects in all three systems (**Figure 3D, E**, and **SI Figure S21**). Further analysis found Gdm^+^ ions to accumulate at the inner nanopore surface, particularly near the termini of the α-hemolysin stem, see **Figure 3F** (top) and **SI Figure S22**. In the same regions, individual Gdm^+^ ions were observed to remain bound to the nanopore surface for considerable (> 10 ns) intervals of time, see **Figure 3F** (bottom). In all systems, the local concentrations of ionic species were found to satisfy the local electroneutrality condition (**SI Figure S23**). However, in the GdmCl/KCl system, K^+^ ions were almost excluded from the α-hemolysin stem (**SI Figure S23**), which explains their negligible current. Thus, binding of Gdm^+^ ions to the inner nanopore surface renders the surface positively charged. That surface charge is compensated by much more mobile chloride ions that carry most of the ionic current and produce a strong electroosmotic effect.

The electroosmotic effect produced by Gdm^+^ binding produced a small yet measurable net transport of the unfolded protein fragments through the nanopore. We next computed the number of residues translocated through the nanopore constriction as a function of simulation time (**Figure 3G** and **SI Figure S24**). To exclude the effect of peptide chain shrinking or stretching, we identified the parts of the simulation trajectories where the number of peptide residues within the α-hemolysin stem remained approximately constant (**SI Figure S25**). Averaged over such constant-density trajectory fragments, the peptides were found to move with the average rate of 1.0+/−0.8 and 0.8+/−0.5 residues/µs for pure and mixed GdmCl electrolytes, respectively, and 0.1+/−0.4 residues/microsecond for pure KCl (**Figure 3H**).

## Protein-specific current signals

In order to extract an “average shape” of ionic current signals produced by the translocation of C-tagged MBP (MBP-D10), N-tagged MBP (D10-MBP), and C-tagged MBP dimer (diMBP-D10), their barycenters (Fréchet means) were computed using the Soft Dynamic Time Warping (SDTW) metric^53^. The result of the barycenter computation is a smooth curve, representing the centroid, or the “essence” of the translocation event shapes in the dataset. The barycenters of each of the variant datasets are shown as colored line in **Figure 4A**, superimposed on resampled events shown in the background (black). The event selection criteria (dwell time and current maximum ranges), the number of passing events, and computation parameters are listed in **Table S4**. In short, we screened events with a narrow range of dwell times near the mean of each variant’s distribution (3-5 ms for MBP-D10 and D10-MBP, and 6-9 ms for diMBP-D10), which later allowed the SDTW algorithm to compute a clear and distinct signature shape for each variant. The current traces of the selected events were then resampled via interpolation to a segment count proportional to the average duration (300 points for MBP-D10 and D10-MBP, and 500 points for diMBP-D10). The resampling reduces the effect of dwell-time variation on this SDTW computation. The resulting barycenter curves show a trend of how the events of each protein type tend to progress, on average. Looking at the positioning of the local maxima and minima, as well as the slope in the middle of the barycenter, we note that MBP-D10 and D10-MBP events show matching opposite trends (**Figure 4A**, left and middle panel). Whether due to the proteins remaining secondary structure in the pore, or purely due to sequence variation, the opposing current blockage trends are a strong indication of directional protein translocation. Further, the skewed “W” shape of diMBP-D10 barycenter and the position of its local minima and maxima appears similar to the MBP-D10 barycenter but repeated twice, especially the “bumps” in the beginning and the middle of the barycenter (**Figure 4A**, right panel).

**Figure 4.**
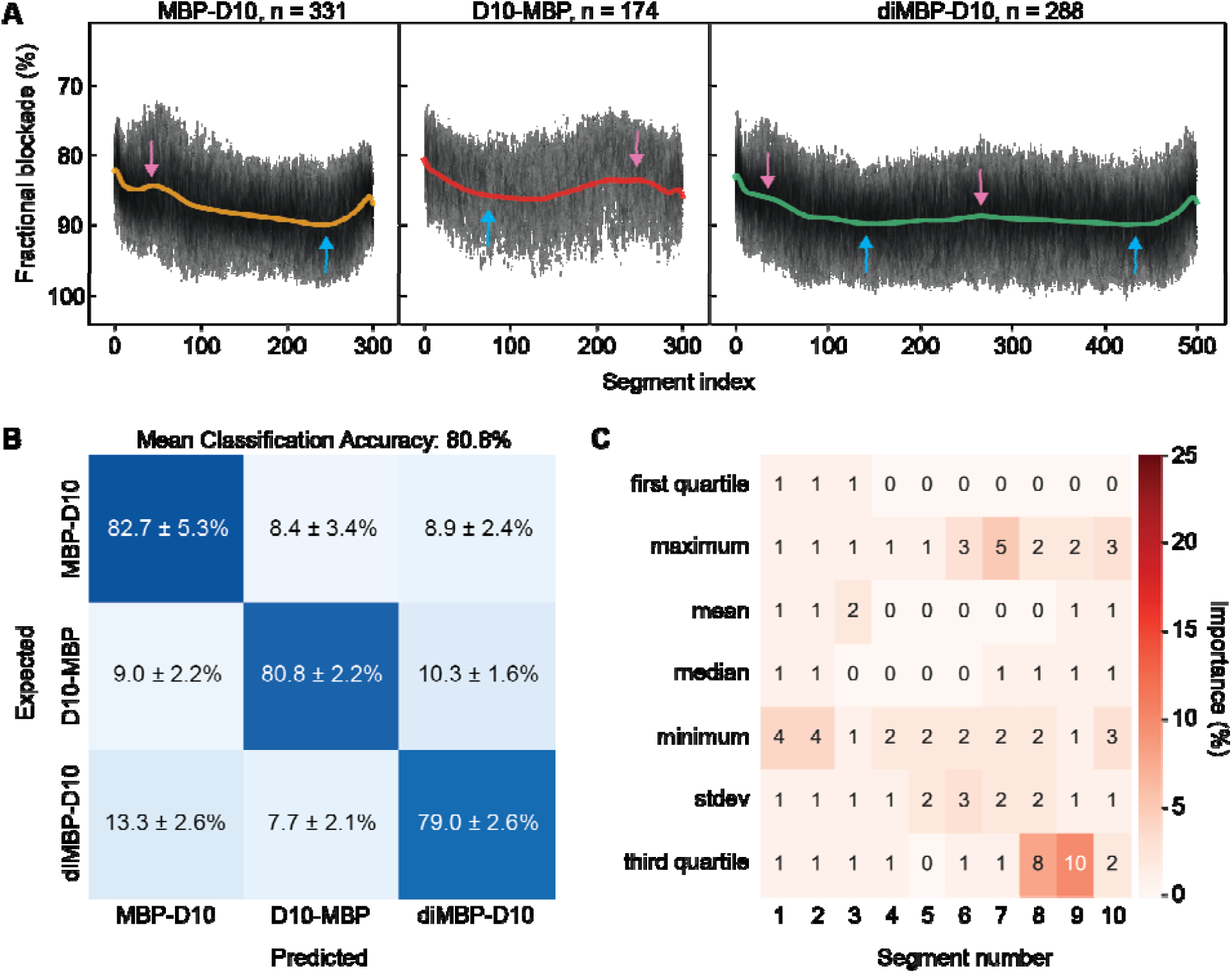
Unidirectional translocation and discrimination of MBP variants. **A)** Soft-DTW barycenters (solid curves) showing resampled events of MBP-D10 (left), D10-MBP (middle) and diMBP-D10 (right), with their respective events superimposed in the background (black). The positions of identified maxima (pin arrows) and minima (blue arrows) are shown. **B)** Three-way confusion matrix representing the mean classification accuracy of a multiclass gradient boosting classifier (GBC) trained to discriminate MBP-D10, D10-MBP, and diMBP-D10 (mean+/−standard deviation of 9 GBC models, each trained and tested on 80% of the data from a reshuffled dataframe, shown in **SI Figure S27**). **C)** A heatmap showing the relative importance of each feature used to generate the multiclass GBC model for discrimination of MBP variant (all 70 features sum to 100%). Each column of the heatmap represents a segment index of the event, and each row represents a statistical parameter extracted from every segment (refer to **SI Figure S27** for further details on features). All experiments were performed in 1.0 M KCl, 2.0 M GdmCl, 10 mM Tris, pH 7.5 and under a 175 mV bias applied to the *trans* chamber.

As the haziness of the resampled events in the background show (**Figure 4A**), transport of a given molecule could have velocity variations at different depths of translocation. Analogous to velocity variations in DNA and RNA ratcheting enzymes in sequencing, it could be possible to computationally eliminate the effect of the velocity variation in protein translocation using a neural network analysis pipeline. However, the highest translocation velocities must not exceed the bandwidth capacity of the nanopore instrument to avoid loss of signal. Because the α-hemolysin pore used does not have sufficient amino acid resolution, we have not attempted any further investigations of the velocity changes in our protein translocation data.

To investigate whether the signal properties from single events comprise sufficient information to discriminate among different protein variants and protein classes, we employed supervised machine learning (ML). A gradient boosting classifier (GBC) was trained and tested for discrimination among MBP variants (MBP-D10, D10-MBP, and diMBP-D10) and distinguishing MBP-D10 from GFP-D10 in a binary mixture. Both GBC models were generated and evaluated with features extracted from labeled translocation events recorded one protein type at time, where 80% of the data were used to train the model and the remaining 20% were withheld for testing the model’s accuracy (see **SI Supplementary Note 4**). While on a population-wide analysis, dwell time alone could inform on the ratio of protein types in a binary mixture (see **SI Figure S26)**, single-molecule identification requires more information from the signal. Hence, blockade-based features from translocation events (dwell time ranging between 300 µs and 20 ms) were used to train and test both GBC models. Specifically, each event was divided into ten segments of equal length and seven statistical parameters from every segment were extracted, creating a feature space comprised of 70 dimensions for GBC model input (see **SI, Figure S27** and **Supplementary Note 4**).

As confusion matrices in **Figure 4B** and **Figure 5E** show, mean classification accuracies of 80.8% and 89.6% were achieved with GBC models trained for three-way classification of MBP variants and two-way classification of MBP-D10 and GFP-D10, respectively. Each confusion matrix is an average of nine reshuffled combinations of samples allocated into the training set and testing set (see **SI, Supplementary Note 4** and **Figures S28** and **S29**). We investigated the relative importance of each feature for GBC prediction in each case (**Figure 4C** and **Figure 5F**). We found the third quartile value in the 8^th^ and 9^th^ event segments to have the highest predictive power for discrimination of MBP variants. This suggests that there is a distinguishable blockade current near the end of MBP-D10, D10-MBP and diMBP-D10 translocation. In contrast, the current standard deviation of the 2^nd^, 3^rd^, and 4^th^ event segments had the greatest influence on GBC classification of MBP-D10 and GFP-D10. To support this result, we observed the greatest difference in local volume standard deviation to occur around these specific segments (see **SI, Figure S30** and **S31**).

**Figure 5.**
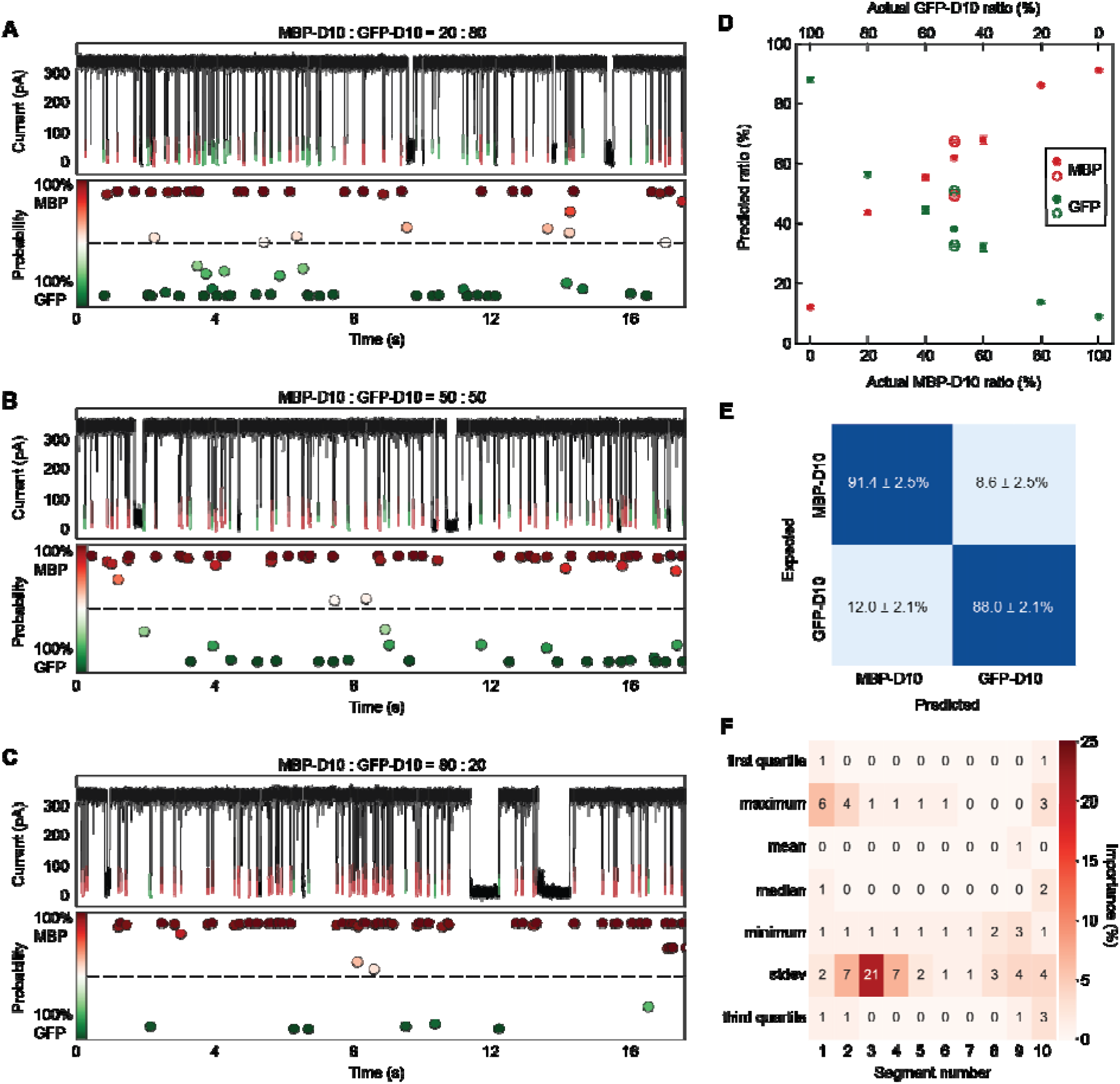
Single-molecule fingerprinting of full-length MBP-D10 and GFP-D10. **A-C)** Post-training GBC classification results for unlabeled MBP-D10 (red) and GFP-D10 (green) mixture experiments at 20:80 **(A)**, 50:50 **(B)** and 80:20 **(C)** ratios with a probability classification estimate associated with each translocation event. **D)** Percentage of MBP-D10 (red) to GFP-D10 (green) predicted by GBC model (shown on the y-axis) when applied to different MBP-D10:GFP-D10 ratio experiments. Each marker is the result of 9 GBC models, each fit with 80% of the training data from a reshuffled dataframe containing features from pure MBP-D10 and GFP-D10 experiments (mean+/− standard deviation). Experiments conducted on different days at 50:50 ratio are shown as hollow markers, highlighting experimental variability. Refer to **SI Table S6** for sample size of events parsed for each ratio experiment. **E)** Two-way confusion matrix representing the mean classification accuracy of a GBC model trained to discriminate MBP-D10 and GFP-D10 (mean+/−standard deviation of 9 GBC models, each trained and tested on 80% of the data from a reshuffled dataframe, shown in **SI Figure S28**). **F)** A heatmap showing the relative importance of each feature used to generate the multiclass GBC model for discrimination of MBP-D10 and GFP-D10 (all 70 features sum to 100). Each column of the heatmap represents a segment index of the event, and each row represents a statistical parameter extracted from every segment (refer to **SI Figure S27** for further details on features). All experiments were performed in 1.0 M KCl, 2.0 M GdmCl, 10 mM Tris, pH 7.5 and under a 175 mV bias applied to the *trans* chamber. The combined concentration of MBP-D10 and GFP-D10 for every ratio experiment is 0.70 μM.

To validate the discrimination capability of the trained model, we tested its performance on unlabeled mixture experiments. We note that simply deploying the model on a 50:50 mixture would not provide any useful information. Instead, we applied the trained GBC model on unlabeled events from mixture experiments containing different MBP-D10:GFP-D10 molar ratios and expected the classification ratios to follow the same trend. **Figures 5A-C** show experimental traces after GBC classification for experiments with 20:80 **(A)**, 50:50 **(B)**, and 80:20 **(C)** MBP-D10 (red) to GFP-D10 (green) mixture, with the model’s confidence score for each call labeled below the trace (see **SI, Supplementary Note 4** and **Table S6, S7**). Plotting the ratio of MBP-D10 to GFP-D10 predicted by the model versus the true concentration of MBP-D10 (**Figure 5D**), we observe ~10% error by the model at an actual MBP-D10 ratio of 0%, followed by a trend where the number of MBP-D10 events called by the model increases roughly linearly with the actual concentration of MBP-D10 (as expected), and a saturation at around 90% when the actual ratio of MBP-D10 is 100%. Model calls where MBP-D10 is higher than expected (20%, 40%, and 50%) could be attributed by the higher capture rate observed with MBP-D10 in comparison to GFP-D10 (**SI Figure S18**). Example events from the training sets and mixture classification results are provided in **SI Figure S32**. The total event ratio predicted by the GBC model for 50:50 mixture experiment is in excellent agreement with the respective population size obtained by integrating the mixture experiment’s dwell time histogram after fitting with the 1D drift-diffusion model^54–57^ (see **SI, Supplementary Note 2** and **Figure S19**).

## Conclusions

We have demonstrated here a method for voltage-driven, single-file transport of full-length proteins in their fully denatured state through a biological nanopore. We find that the guanidinium ions that mediate the protein unfolding in preparation for transport also facilitate electroosmotic flow through the pore through their interactions with the hemolysin pore. This electroosmotic flow that can drive an unstructured protein chain through a pore is important not only in the enzyme-free approach we presented here, but may also find use for keeping a protein chain taut at the pore during for enzyme-mediated nanopore-based single-molecule protein sequencing, required for reliable readout of the amino acids in the chain at the pore constriction. For the protein molecules studied here, we find mean transport speeds of ~10 µs/amino acid. On average, this is slow enough to collect several current datapoints for each protein segment in the pore constriction, although the actual measurement time resolution is noise-limited and further work is needed to reduce this noise and achieve higher bandwidths in protein pore measurements. We also find that the orientation and number of repeat units of a protein can be distinguished based on the current signatures (i.e., N-to-C vs. C-to-N transport and monomer vs. dimer). Further, in binary mixture experiments consisting of tewo proteins we show that machine-learning analysis can distinguish between two proteins based on their single-molecule electrical signatures with ~90% accuracies. Analysis of native proteins by can also be accomplished after end-tagging with DNA at the N-terminus, although efficiencies for this end-tagging reaction were low (~6%) and require optimization.

We note here several improvements that need to be made for using GdmCl-assisted unfolding and translocation through nanopores. First, one needs to develop a facile route to high-efficiency protein functionalization at either the N- or C- to enable analyte capture and threading. Next, electronic noise should be reduced so that high-bandwidth measurements can be conducted with GdmCl-mediated protein transport through a higher-resolution pore (such as MspA, which maintains its functionality in 2 M GdmCl^58^), to increase the resolution of protein fingerprinting. Finally, although we use GdmCl for threading/stretching a protein in the pore, we note that combining this mode with enzyme-driven protein motion is possible, for example, by using an asymmetric buffer in which: 1) protein unfolding is mediated in a denaturant chamber, 2) electroosmotic forces are used to keep the protein taut at the pore, and 3) enzyme-mediated motion on another chamber of the pore is used to move the protein through the pore in a discrete manner. Other improvements such as changing the electrolyte type^59^, pore variant^60–62^, and voltage waveform, can be made to further enhance the ability of our method to obtain more unique and unequivocal fingerprint signatures from full-length proteins.

## Methods

### Polymer bilayer painting and nanopore measurement

The 100 µm SU-8 wedge-on-pillar aperture supported by a 500 µm-thick Si chip with a square open window^39^ was mounted on our custom-designed fluidic cell, sealing properly to separate *cis* and *trans* chambers. Both sides of the aperture were pretreated with 4 mg/ml poly(1,2-butadiene)-b-poly(ethylene oxide) (PBD_11_– PEO_8_) block-copolymer (Polymer Source) dissolved in hexane to coat the aperture with a dry and thin polymer layer. The *cis* and *trans* chambers were filled with GdmCl electrolyte (all contain 1 M KCl, 10 mM Tris, pH 7.5), and a pair of Ag/AgCl electrodes were immersed in the electrolyte and connected to an Axon 200B patch-clamp amplifier. The polymer membrane was painted across the aperture using 8 mg/ml polymer dissolved in decane. At least 60 mins waiting time were required until the polymer membrane thinned to a capacitance value of 60 – 80 pF. After verification of bilayer formation, 0.5 µl of 50 µg/ml α-hemolysin (Sigma-Aldrich) was added to the *cis* chamber, and an ion conductance jump marked single pore insertion. A denatured protein sample (incubated in GdmCl buffer before use) was added to the *cis* chamber and mixed gently by pipetting. Current signals were low-pass filtered at 100 kHz using the Axopatch setting and digitized at 16-bits and 250 kHz sampling rate using a National Instruments Data Acquisition card and custom LabVIEW-based software that records and saves all raw current data and acquisition settings. In further data analysis digital low-pass filtering was used to further reduce the noise.

### Cloning of the GFP and MBP constructs

All primers (Eurofins MWG Operon) used in this study are listed in **Table S1**. The N-terminal 10-aspartate MBP (D10-MBP), C-terminal 10-aspartate MBP (MBP-D10), and C-terminal 10-aspartate GFP (GFP-D10) constructs were obtained by mutagenesis polymerase chain reaction (PCR) using pT7-MBP or pRSETB-GFP as the template plasmid. The PCR reaction mixtures were subjected to DpnI digestion for 3 h at 37 °C to degrade the template plasmids. The digested samples were then transformed into chemically competent *E. coli* DH5α cells. The desired mutant plasmids were isolated from colonies and verified by DNA sequencing.

The C-terminal MBP-D10 dimer construct (diMBP-D10) was generated as follows: the first mutagenesis PCR was performed using pT7-hisMBP as the template to remove the stop codon and add a flexible linker GGSG to the C-terminus of the MBP gene. The PCR products were digested with DpnI and transformed into *E. coli* DH5α cells resulted in a plasmid pT7-hisMBPggsg containing the hindIII and Sfbl restriction sites right after the GGSG linker gene. The second PCR was performed with pT7-MBP as the template to introduce HindIII and Sfbl cutting sites at the two ends of the MBP gene and add a D10 at the c-terminal to the MBP fragment. The PCR products and the plasmid pT7-hisMBPgsgg were digested with HindIII and Sfbl and ligated by T4 ligase. The ligated products were transformed into chemically competent *E. coli* DH5α cells. The mutant plasmid pT7-diMBP-D10 was verified by enzyme digestion and DNA sequencing.

### Expression and purification of GFP and MBP proteins

GFP and MBP protein variants (**Table S2**) were expressed and purified using similar protocols. Briefly, plasmids were transformed into chemically competent BL21(DE3) *E. coli* cells. The cells were grown in 1 L of LB medium at 37 °C until the OD600 reached 0.6 and induced with 0.5 mM isopropyl β-D-1-thiogalactopyranoside. The temperature was then decreased to 16°C for overnight expression. Cells were harvested by centrifugation at 13000 RPM for 25 min. The cell pellets were used for protein purification or frozen at −20 °C for future use. Cells were resuspended in 50 ml of 50 mM Tris-HCl (pH 8.0), 150 mM NaCl buffer, and lysed via sonication to purify proteins. The lysate was centrifuged at 13000 RPM for 25 min. The supernatant was filtered through a 0.22 µm syringe filter (CELLTREAT Scientific Products) and then loaded to a Ni-NTA affinity column (ThermoFisher scientific) equilibrated with buffer 50mM Tris-HCl (pH 8.0), 150 mM NaCl. MBP-D10, D10-MBP and diMBP-D10 were eluted in buffer 50 mM Tris-HCl (pH 8.0), 150 mM NaCl, 150 mM imidazole. GFP-D10 was eluted in buffer 50 mM Tris-HCl (pH 8.0), 150mM NaCl, 20 mM imidazole. After Ni-NTA chromatography, MBP-D10, D10-MBP, and GFP-D10 exhibited more than 95% purity on SDS-PAGE, while the eluted diMBP-D10 fraction contained multiple low-molecular impurity bands. The eluted samples were run on a preparative 12 % SDS-PAGE to remove these impurity proteins. The band containing the full-length diMBP-D10 was cut out, and the protein was extracted from the gel with buffer 50mM Tris-HCl (pH8.0), 8M urea by incubating the gel and the extraction buffer at room temperature overnight. The supernatant containing the protein was collected by centrifuging the samples at 13000 RPM for 30 min. Protein concentrations of all samples were determined by A280 with Nanodrop and stored at −80 °C for future use.

### MD simulation

All MD simulations were performed using the molecular dynamics program NAMD2^63^, a 2 femtosecond integration timestep, periodic boundary conditions, CHARMM36^64^ force field, and a custom non-bonded fix (NBFIX) corrections for K, Cl, and Gdm ions^65^. SETTLE algorithm^66^ was used to maintain covalent bonds to hydrogen atoms in water molecules, whereas RATTLE algorithm^67^ maintained all other covalent bonds involving hydrogens. The particle-mesh Ewald^68^ method was employed to compute long-range electrostatic interactions over a 1.2 Å grid. All van der Waals and short-range electrostatic interactions were evaluated every time step using a cutoff of 12 Å and a switching distance of 10 Å; Full electrostatics were evaluated every second-time step.

The all-atom models of α-hemolysin suspended in a lipid bilayer membrane were built using CHARMM-GUI^69^. The initial structural model of α-hemolysin was taken from the Protein Data Bank (PDB ID: 7AHL)^70^. After adding missing atoms and aligning the primary principal axis of the protein with the *z*-axis, the protein structure was merged with a 15×15 nm^2^ patch of a pre-equilibrated 1-palmitoyl-2-oleoyl-sn-glycero-3-phosphocholine (POPC) lipid bilayer. The protein-lipid complex was then solvated in a rectangular volume of ~78,500 pre-equilibrated TIP3P water molecules^71^. Gdm^+^, K^+^, and Cl^−^ ions were added at random positions corresponding to target ionic concentrations. Additional charges were introduced to neutralize the system. Each final system was 15×15×18 nm^3^ in volume and contained approximately 300,000 atoms. Upon assembly, the systems were initially equilibrated using the default CHARMM-GUI’s protocol. Specifically, the systems were subjected to energy minimization for 10,000 steps using the conjugate gradient method. Next, lipid tails and protein side chains were relaxed in a 2.5 ns pre-equilibration simulation that was run while restraining the protein backbones and lipid head groups. This step was followed by a 25 ns simulation in the NPT (constant number of particles, pressure, and temperature) ensemble using the Nosé-Hoover Langevin piston pressure control^72^. In all simulations, the temperature was maintained at 298.15 K by coupling all non-hydrogen atoms to a Langevin thermostat with a damping constant of 1 ps^−1^.

The atomic coordinates of the maltose-binding protein (MBP) were obtained from the Protein Data Bank (entry 1JW4)^73^. The missing hydrogen atoms were added using the psfgen plugin of VMD^74^. The protonation state of each titratable residue was determined using PROPKA^75^ according to the experimental pH conditions (7.5 pH). Next, the protein was split into seven peptide fragments producing six 53-residue and one 52-residue fragments. The N-terminal of each peptide was terminated with a neutral acetyl group (ACE patch), whereas the C-terminal was terminated with an N-methyl group (CT3 patch). Each peptide was stretched using constant velocity SMD in vacuum, followed by 5 ns equilibration in a 1.5 M GdmCl solution. During the 150 ps SMD run, the C-terminal of the peptide was kept fixed. At the same time, the N-terminal was coupled to a dummy particle utilizing a harmonic potential (*k*_spring_ = 7 kcal/(mol Å^2^)), and the dummy particle was pulled with a constant velocity of 1 Å/ps. At the end of the equilibration step, each peptide fragment had a contour length of approximately 167 Å, ~3.16 Å per residue. Next, we used the phantom-pore method^50^ to convert the geometrical shape of the α-hemolysin nanopore to a mathematical surface. To fit the stretched peptide into the α-hemolysin pore, the phantom pore surface was initially made to represent a nanopore that was 1.4 times wider than the pore of α-hemolysin. During a 2 ns simulation, the phantom pore was gradually shrunk to match the shape of the α-hemolysin nanopore. At the same time, all atoms of the peptide and all ions laying outside the potential were pushed toward the nanopore center using a constant 50 pN force. At the end of the simulation, each peptide fragment and all guanidinium ions residing within 3 Å of any peptide atom were placed inside the pre-equilibrated α-hemolysin system having the peptide’s backbone approximately aligned with the nanopore axis. Before the production runs, each system was equilibrated for 10 ns in the NPT ensemble at 1.0 bar, and 298.15 K with all Cα atoms of the α-hemolysin protein restrained to the crystallographic coordinates.

All production simulations were carried out in the constant number of particles, volume, and temperature ensemble (NVT) under a constant external electric field applied normal to the membrane, producing a ±200 mV transmembrane bias. To maintain the nanopore’s structural integrity, all protein’s C_α_ atoms were restrained to exact coordinates as in the last frame of the equilibration trajectory using harmonic potentials with spring constants of 1 kcal/(mol Å^2^). The ionic currents were calculated as described previously^52^. The quantify protein translocation, we defined the number of residues translocated as the number of non-hydrogen backbone atoms passing below the α-hemolysin constriction divided by the total number of non-hydrogen backbone atoms in one residue. The constriction’s z-coordinate was defined by the center of mass of the backbone atoms of residues 111, 113, and 147. The concentration profile and guanidinium binding analyses were carried out using in-house VMD scripts. All MD trajectories were visualized using VMD^74^.

## Data analysis and code availability

All data parsing (excluding DTW and GBC) were performed using the Pyth-ion package (https://github.com/wanunulab/Pyth-Ion) and figures were generated using Igor. For analysis, the raw 100 kHz nanopore current data was further low-pass filtered to 10 kHz using the low-pass filter function in Pyth-ion. DTW and GBC analyses were conducted via python scripts written and documented in Jupyter Notebooks, tslearn (v0.5.2)^76^, SciKit-Learn (v1.0.2)^77^, and a modified version of the PyPore^78^ nanopore data analysis library. The Jupyter notebook and associated files are available on GitHub (https://github.com/wanunulab/protein-gd). A detailed description of the DTW and GBC analyses is provided in **SI Section 6**.

## Supporting Information Available

Data analysis details, example datasets, and methods description. All data used in this manuscript are available for download at https://figshare.com/s/5cd39ee415c62a316a6f.

## Author Contributions

L.Y. and M.W. conceived the project and designed the experiments. L.Y. developed the experimental protocol, carried out the experiments, and analyzed the protein translocation data. X.K. and L.Y. fabricated the SU-8 wedge-on-pillar chips used in the experiments. F.L. and M.C. prepared and purified the protein samples. B.M. and A.A. designed and conducted the MD simulations. A.M. and A.F. performed the data analysis described in “protein specific current signals” (SDTW and GBC) and wrote the respective sections of the manuscript. J.C.F performed the DNA-protein conjugation experiments. L.Y., B.M., A.A, and M.W. wrote the first draft of the manuscript, and all authors commented and edited it.

## Acknowledgment

We thank Caroline McCormick for assistance with editing the manuscript for clarity, and Dr. Nikolai Slavov for helpful discussions regarding protein sequencing. We acknowledge funding from the National Institutes of Health: HG0011087 (MW); GM115442 (MC), and the National Science Foundation: PHY-1430124 (AA). The supercomputer time was provided through the XSEDE allocation grant (MCA05S028) and the Leadership Resource Allocation MCB20012 on Frontera of the Texas Advanced Computing Center.

