## Supplementary material for "Unidirectional Single-File Transport of Full-Length Proteins Through a Nanopore": SI

### Table of Contents

Section 1: Information on proteins in this study

a. **Table S1:** List of primers for cloning GFP and MBP constructs
b. **Table S2:** Amino acid sequences of the protein constructs in this study
c. **Figure S1:** SDS-PAGE analysis of MBP N-terminus oligonucleotide labeling
d. **Supplementary Note 1:** N-terminal Oligonucleotide labeling of MBP
e. **Figure S2:** An example current trace and fractional current blockade vs. dwell time scatter plot for T20-MBP
f. **Figure S3:** 12% SDS-PAGE analysis of MBP and GFP proteins

Section 2: Extra data and analysis for MBP-D10

a. **Figure S4:** Effect of [GdmCl] on MBP-D10 interactions with α-hemolysin
b. **Figure S5:** Fractional current blockade vs. dwell time scatter plots at 11 different voltages for MBP-D10 in 1.0 M GdmCl
c. **Figure S6:** Current vs. time traces at 11 different voltages for MBP-D10 in 1.0 M GdmCl
d. **Figure S7:** Fractional current blockade vs. dwell time scatter plots at 10 different voltages for MBP-D10 in 1.5 M GdmCl.
e. **Figure S8:** Current vs. time traces at 10 different voltages for MBP-D10 in 1.5 M GdmCl
f. **Figure S9:** Fractional current blockade vs. dwell time scatter plots at 10 different voltages for MBP-D10 in 2.0 M GdmCl
g. **Figure S10:** Current vs. time traces at 10 different voltages for MBP-D10 in 2.0 M GdmCl

Section 3: Extra data and analysis for diMBP-D10

a. **Figure S11:** Comparison of dwell time histograms between MBP-D10 and diMBP-D10
b. **Figure S12:** Mean dwell time vs. voltage for the $P_F$ populations
c. **Figure S13:** Fractional current blockade vs. dwell time scatter plots at 10 different voltages for diMBP-D10 in 1.5 M GdmCl
d. **Figure S14:** Current vs. time traces at 10 different voltages for diMBP-D10 in 1.5 M GdmCl

e. **Figure S15:** Fractional current blockade vs. dwell time scatter plots and current vs. time traces at 3 different voltages for diMBP-D10 in 2.0 M

Section 4: Other experimental data

a. **Figure S16:** Current vs. time trace for MBP-D10 without GdmCl at V = 175 mV
b. **Figure S17:** $P_L$ percentage vs. voltage plots for MBP-D10 and diMBP-D10 in different GdmCl concentrations
c. **Figure S18:** Concentration-normalized capture rates as a function of voltage for MBP-D10 and diMBP-D10 in different GdmCl concentrations
d. **Figure S19:** Dwell time histograms for pure GFP-D10, pure MBP-D10 and mixture of them with fitting lines based on 1D drift-diffusion model
e. **Supplementary note 2:** Estimate of the event ratio in the MBP-D10/GFP-D10 mixture

Section 5: MD simulation

a. **Table S3:** Fragmentation of the maltose-binding protein (MBP) into seven peptides for MD simulations.
b. **Figure S20:** MD simulation of α-hemolysin systems containing fragments of MBP.
c. **Figure S21:** Simulated electro-osmotic flow in α-hemolysin systems containing fragments of MBP
d. **Figure S22:** Gdm binding to the inner surface of α-hemolysin in MD simulations of the α-hemolysin systems containing MBP fragments
e. **Figure S23:** Local concentration of ions along the transmembrane nanopore observed in MD simulations of the α-hemolysin systems containing fragments of MBP
f. **Figure S24:** Simulated transport of MBP fragments
g. **Figure S25:** Number of residues confined within the stem region of α-hemolysin

Section 6: Machine learning analysis

a. **Table S4:** Selection and processing parameters for translocation events used in the directionality analysis via Soft-DTW
b. **Figure S26:** Fractional current blockade vs. dwell time scatter plots and corresponding dwell time histograms for different GFP-D10:MBP-D10 concentration ratios
c. **Table S5:** Sample size of translocation events for each protein type from pure experiments
d. **Table S6:** Sample size of translocation events parsed from MBP-D10:GFP-D10 mixture experiments
e. **Table S7:** Predicted ratios by trained GBC model on unlabeled events parsed from MBP-D10:GFP-D10 mixture experiments
f. **Figure S27:** Graphical representation of the 7 statistical parameters extracted from each segment (S1 to S10) of a full MBP-D10 translocation event
g. **Figure S28:** MBP-D10 vs. D10-MBP vs. diMBP-D10 all iterations of training and testing
h. **Figure S29:** MBP-D10 vs. GFP-D10 all iterations of training and testing
i. **Figure S30:** Separation of top 3 weighted features for MBP-D10 and GFP-D10
j. **Figure S31:** Stdev. of volume after dividing MBP-D10/GFP-D10 into 5 segments
k. **Figure S32:** Comparison between event traces for pure MBP-D10 or GFP-D10 and event traces classified by trained GBC model
l. **Figure S33:** Soft-DTW barycenter selected via different dwell-time restriction criteria
m. **Supplementary Note 3:** Soft-DTW Analysis
n. **Supplementary Note 4:** Gradient Boosting Classifiers Analysis

### Section 1: Information on proteins in this study

**Table S1.** List of primers for cloning GFP and MBP constructs

| Desired construct | Template plasmid | Primers |
|---|---|---|
| N-terminal 10-aspartate MBP (D10-MBP) | pT7-MBPhis | Forward: 5'-GATGATGACGACGATGATGATAAAATCGAAGAAGGTAAAC-3' |
| | | Reverse: 5'-ATCGTCGTCATCATCGTCATCATCCATATGTATATCTCCTTC-3' |
| C-terminal 10-aspartate MBP (MBP-D10) | pT7-hisMBP | Forward: 5'-GATGATGACGACGATGATGATTAAAAGCTTGGATCC-3' |
| | | Reverse: 5'-ATCGTCGTCATCATCGTCATCATCCTTGGTGATACGAGTC-3' |
| C-terminal 10-aspartate GFP (GFP-D10) | pRSETB-GFP | Forward: 5'-CGATGATGACGACGATGATGATTAAAAGCTTGATCCGGCTGCTAACAAAGCCCGAAAG-3' |
| | | Reverse: 5'-CATCGTCGTCATCATCGTCATCATCTTTGTATAGTTCATCCATG-3' |
| MBP-D10 with HindIII and SbfI cut sites | pT7-hisMBP-D10 | Forward: 5'-ATGATGAAGCTTAAAATCGAAGAAGGTAAAC-3' |
| | | Reverse: 5'-TTCGGACCTGCAGGTTAATCATCATCGTCG-3' |
| MPB with GS-linker at C-terminus, no stop codon | pT7-hisMBP | Forward: 5'-AGGGTAGTAAGCTTGATCCGGCTGCTAACAAAG-3' |
| | | Reverse: 5'-TCAAGCTTACTACCCTTGGTGATACGAGTCTG-3' |

**Table S2.** Amino acid sequences of the protein constructs in this study

| |
|---|
| **N-terminal D10 MBP (D10-MBP)**<br>MDDDDDDDDDDKIEEGKLVIWINGDKGYNGLAEVGKKFEKDTGIKVTVEHPDKLEEKFPQVA<br>ATGDGPDIIFWAHDRFGGYAQSGLLAEITPDKAFQDKLYPFTWDAVRYNGKLIAYPIAVEALS<br>LIYNKDLLPNPPKTWEEIPALDKELKAKGKSALMFNLQEPYFTWPLIAADGGYAFKYENGKYD<br>IKDVGVDNAGAKAGLTFLVDLIKNKHMNADTDYSIAEAAFNKGETAMTINGPWAWSNIDTSKV<br>NYGVTVLPTFKGQPSKPFVGVLSAGINAASPNKELAKEFLENYLLTDEGLEAVNKDKPLGAV<br>ALKSYEEELAKDPRIAATMENAQKGEIMPNIPQMSAFWYAVRTAVINAASGRQTVDEALKDA<br>QTRITKHHHHHHHH |
| **C-terminal D10 MBP (MBP-D10)**<br>MHHHHHHHHKIEEGKLVIWINGDKGYNGLAEVGKKFEKDTGIKVTVEHPDKLEEKFPQVAAT<br>GDGPDIIFWAHDRFGGYAQSGLLAEITPDKAFQDKLYPFTWDAVRYNGKLIAYPIAVEALSLIY<br>NKDLLPNPPKTWEEIPALDKELKAKGKSALMFNLQEPYFTWPLIAADGGYAFKYENGKYDIK<br>DVGVDNAGAKAGLTFLVDLIKNKHMNADTDYSIAEAAFNKGETAMTINGPWAWSNIDTSKVN<br>YGVTVLPTFKGQPSKPFVGVLSAGINAASPNKELAKEFLENYLLTDEGLEAVNKDKPLGAVAL<br>KSYEEELAKDPRIAATMENAQKGEIMPNIPQMSAFWYAVRTAVINAASGRQTVDEALKDAQT<br>RITKDDDDDDDDDD |
| **C-terminal D10 dimer MBP (diMBP-D10)**<br>MHHHHHHHHKIEEGKLVIWINGDKGYNGLAEVGKKFEKDTGIKVTVEHPDKLEEKFPQVAAT<br>GDGPDIIFWAHDRFGGYAQSGLLAEITPDKAFQDKLYPFTWDAVRYNGKLIAYPIAVEALSLIY<br>NKDLLPNPPKTWEEIPALDKELKAKGKSALMFNLQEPYFTWPLIAADGGYAFKYENGKYDIK<br>DVGVDNAGAKAGLTFLVDLIKNKHMNADTDYSIAEAAFNKGETAMTINGPWAWSNIDTSKVN<br>YGVTVLPTFKGQPSKPFVGVLSAGINAASPNKELAKEFLENYLLTDEGLEAVNKDKPLGAVAL<br>KSYEEELAKDPRIAATMENAQKGEIMPNIPQMSAFWYAVRTAVINAASGRQTVDEALKDAQT<br>RITKGGSGKIEEGKLVIWINGDKGYNGLAEVGKKFEKDTGIKVTVEHPDKLEEKFPQVAATGD<br>GPDIIFWAHDRFGGYAQSGLLAEITPDKAFQDKLYPFTWDAVRYNGKLIAYPIAVEALSLIYNK<br>DLLPNPPKTWEEIPALDKELKAKGKSALMFNLQEPYFTWPLIAADGGYAFKYENGKYDIKDV<br>GVDNAGAKAGLTFLVDLIKNKHMNADTDYSIAEAAFNKGETAMTINGPWAWSNIDTSKVNYG<br>VTVLPTFKGQPSKPFVGVLSAGINAASPNKELAKEFLENYLLTDEGLEAVNKDKPLGAVALKS<br>YEEELAKDPRIAATMENAQKGEIMPNIPQMSAFWYAVRTAVINAASGRQTVDEALKDAQTRIT<br>KDDDDDDDDDD |
| **C-terminal D10 GFP (GFP-D10)**<br>MHHHHHHSKGEELFTGVVPILVELDGDVNGHKFSVSGEGEGDATYGKLTLKFICTTGKLPVP<br>WPTLVTTFAYGLQCFARYPDHMKQHDFFKSAMPEGYVQERTIFFKDDGNYKTRAEVKFEGD<br>TLVNRIELKGIDFKEDGNILGHKLEYNYNSHNVYIMADKQKNGIKVNFKIRHNIEDGSVQLADH<br>YQQNTPIGDGPVLLPDNHYLSTQSALSKDPNEKRDHMVLLEFVTAAGITHGMDELYKDDDD<br>DDDDDD |

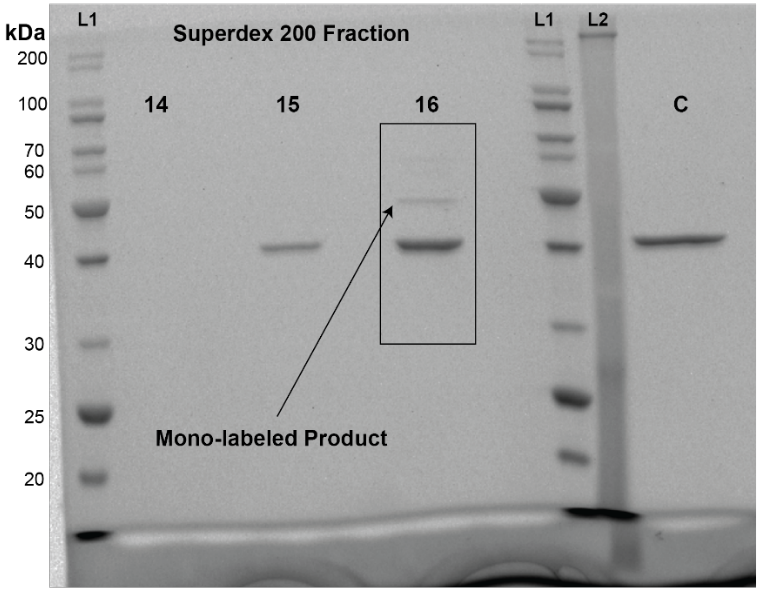


**Figure S1.** SDS-PAGE analysis of MBP N-terminus oligonucleotide labeling. Reaction was purified via Superdex 200 size-exclusion chromatography. Fractions 14-16 appeared to contain labeled product and were thus concentrated via Amicon column. Only fraction 16 contained detectable labeled product (indicated by the black arrow). L1 is unstained broad range protein standard (New England Biolabs), L2 is pre-stained broad range standard (Genescript), and C is unreacted MBP negative control. Gel is a house-made 12% Tris-Glycine gel with stain-free protein visualization accomplished via 2,2,2-Trichloroethanol (TCE).

**Supplementary Note 1: N-terminal Oligonucleotide labeling of MBP**

Twenty nucleotide homothymine functionalized at the 5'-terminus with a benzaldehyde moiety (IDT: /5AhMC2/TTTTTTTTTTTTTTTTTTTT) was used in this procedure. Protein labeling was accomplished via modification of the methods described by Chen et al[1]. Briefly, the reaction was performed in a 4 mL screw-top glass via (Fisher Scientific) with 2.38 nmol MBP (100 mM citric acid buffer pH 6.1), 4.79 nmol benzaldehyde oligo (120 µM in citric acid buffer pH 6.1), 13.47 µmol sodium cyanoborohydride in a 200 µL volume of 100 mM citric acid buffer pH 6.1 (85:15 v/v 100 mM trisodium citrate dihydrate: 100 mM citric acid) at room temperature/15 rpm for 20 hours. Product conversion was assessed via SDS-PAGE band shift. Low conversion to the mono-labeled product was observed, necessitating removal of unreacted labeling reagent. Reaction product was purified via size-exclusion chromatography using a Superdex 200 Increase 10/300 GL column (Cytiva). Fractions containing the product were concentrated using Amicon ultra 0.5 mL 3K mw cutoff (Millipore) under centrifugation at 10,000 rcf. Purity was assessed via SDS-PAGE (**Figure S1**) and protein concentration was determined by A280 via Nanodrop, using extinction coefficient calculated by Benchling GUI.

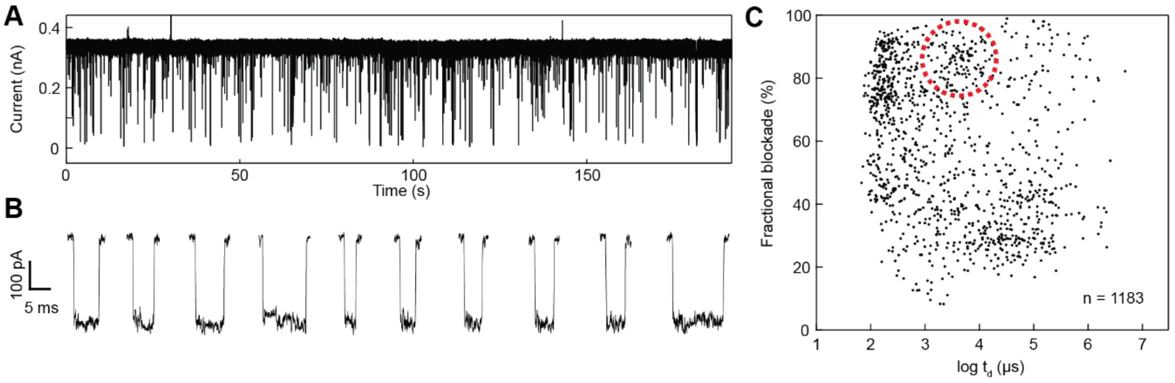


**Figure S2.** An example current trace **(A)** with some example events **(B)**, as well as a fractional current blockade vs. dwell time scatter plot **(C)** for T20 oligonucleotide labeled N-terminus MBP (T20-MBP), where the red dashed circle (including 6.3% of the total events) indicates the same range of translocation events for MBP-D10 in the next section. Experiments were performed in 1.0 M KCl, 2.0 M GdmCl, 10 mM Tris, pH 7.5 with α-hemolysin pore, under a 175 mV bias applied to the *trans* chamber.

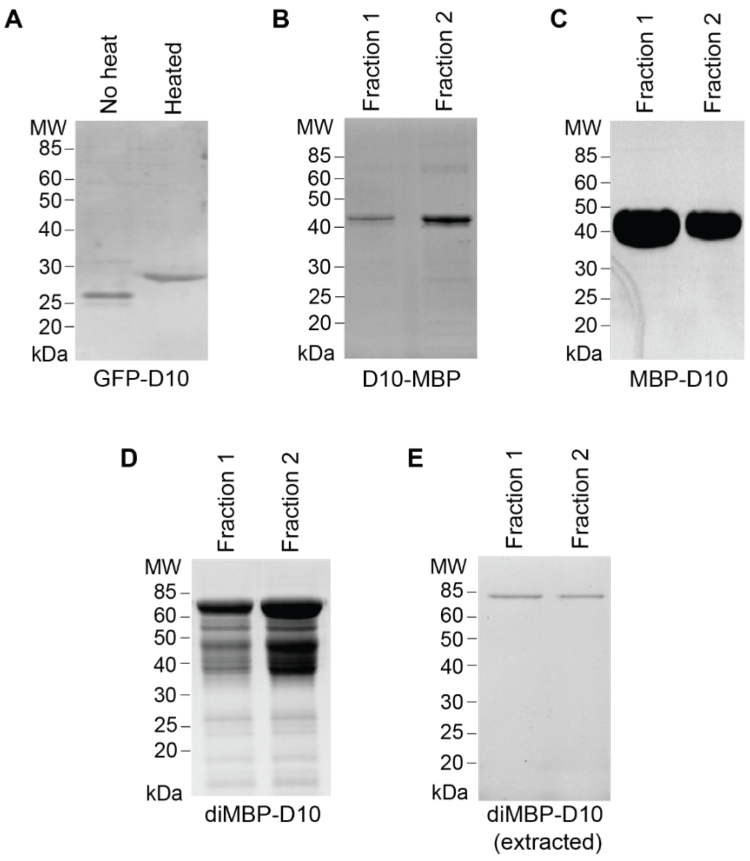


**Figure S3.** 12% SDS-PAGE analysis of MBP and GFP proteins. **A)** Same fraction of C-terminus GFP-D10 after Ni-NTA chromatography purification. Left: no heat; right: heat at 95°C in 50 mM Tris-HCl (pH 8.0), 150 mM NaCl, 20 mM imidazole sample buffer for 15 min. Two fractions of **B)** N-terminus D10-MBP, **C)** C-terminus MBP-D10 and **D)** dimer C-terminus diMBP-D10 after Ni-

NTA chromatography purification. **E)** Two representative fractions of diMBP-D10 after gel extraction purification.

#### Section 2: Extra data and analysis for MBP-D10

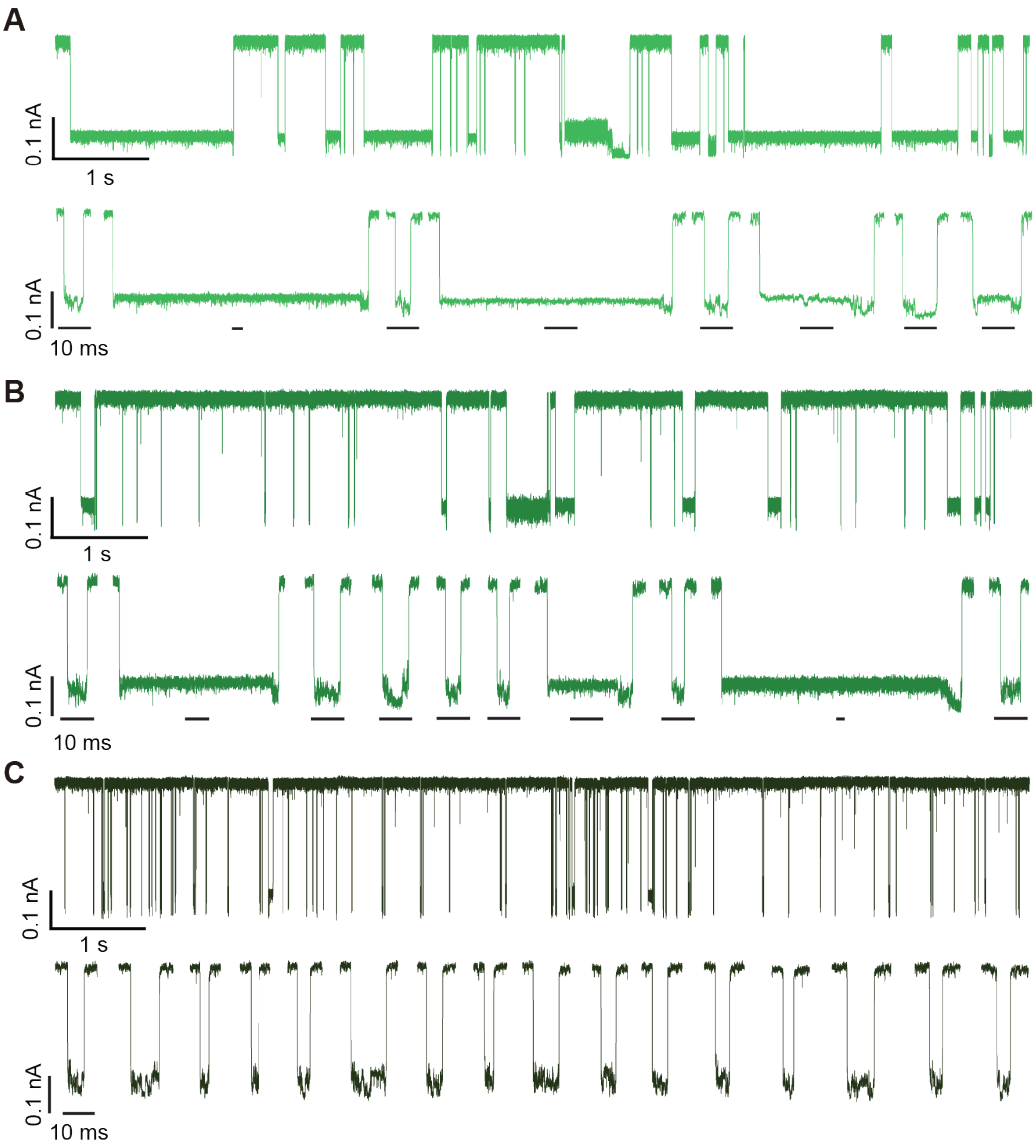


**Figure S4.** Continuous current vs. time traces for MBP-D10 in 1.0 M, 1.5 M and 2.0 M GdmCl, with the open pore currents $I_0$ = 0.260, 0.308, 0.352 nA, respectively. Under each continuous trace we show random events selected at 10 s intervals. All time-scale bars for the selected events are 10 ms. Two populations comprising relatively short (few-ms) and very long (up to 1 s) events are seen in the traces. Further, as the GdmCl concentration increases, the occurrence of long events disappears, and the events are very uniform in duration for the 2.0 M GdmCl case (see **panel C**).

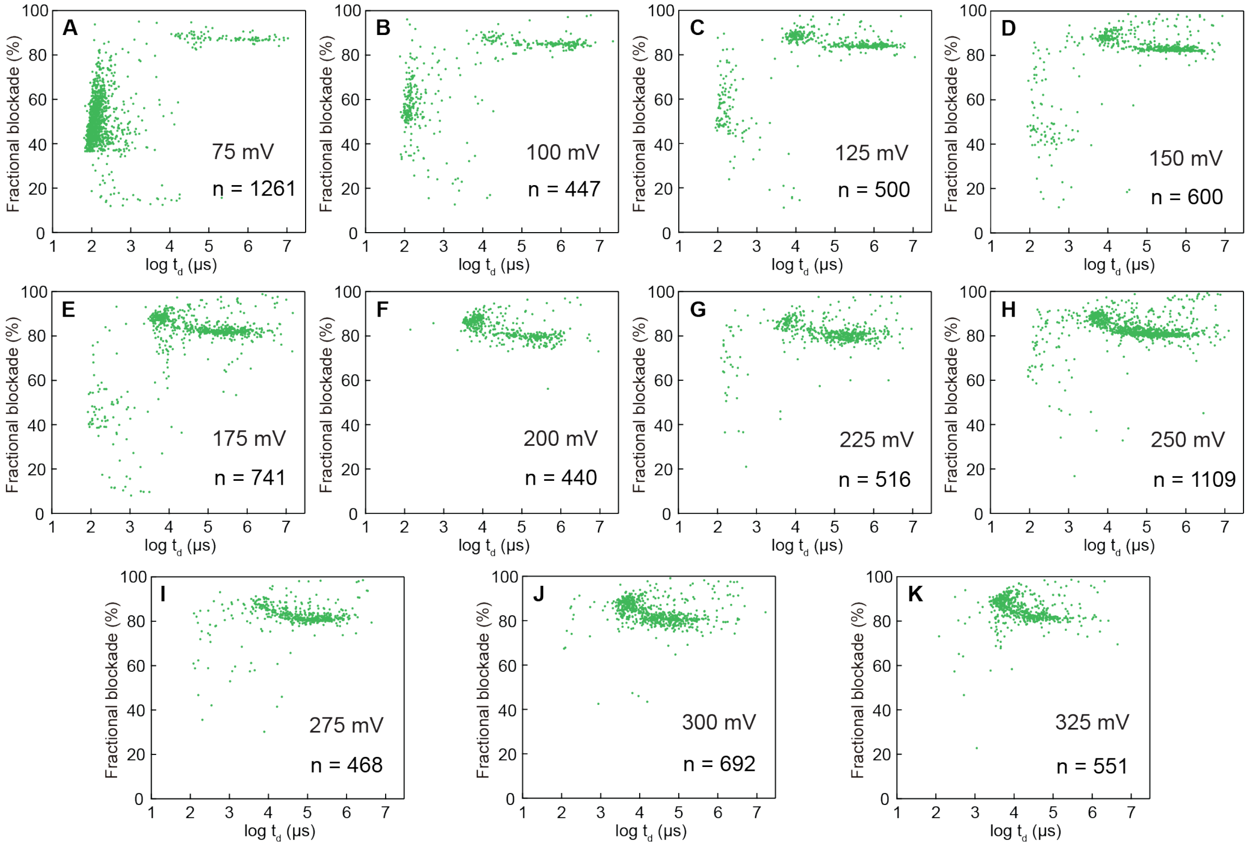


**Figure S5.** Fractional current blockade vs. dwell time scatters for MBP-D10 (0.7 μM) in 1.0 M GdmCl (1 M KCl, 10 mM Tris, pH 7.5), V = 75 – 325 mV.

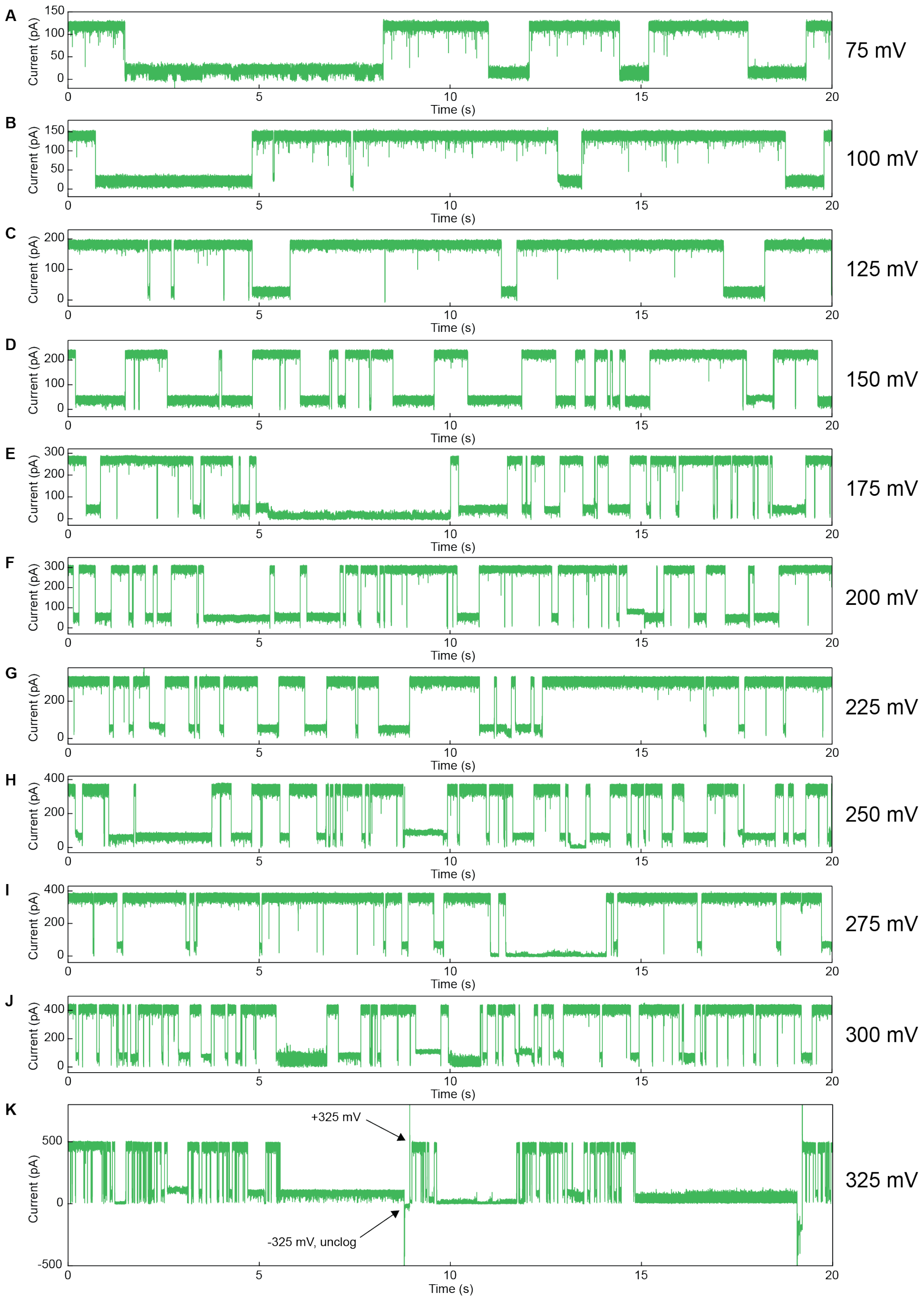


**Figure S6.** Current vs. time traces for MBP-D10 (0.7 µM) in 1.0 M GdmCl (1 M KCl, 10 mM Tris, pH 7.5), V = 75 – 325 mV, where panel K shows that when pore get clogged, switching voltage may help to unclog it.

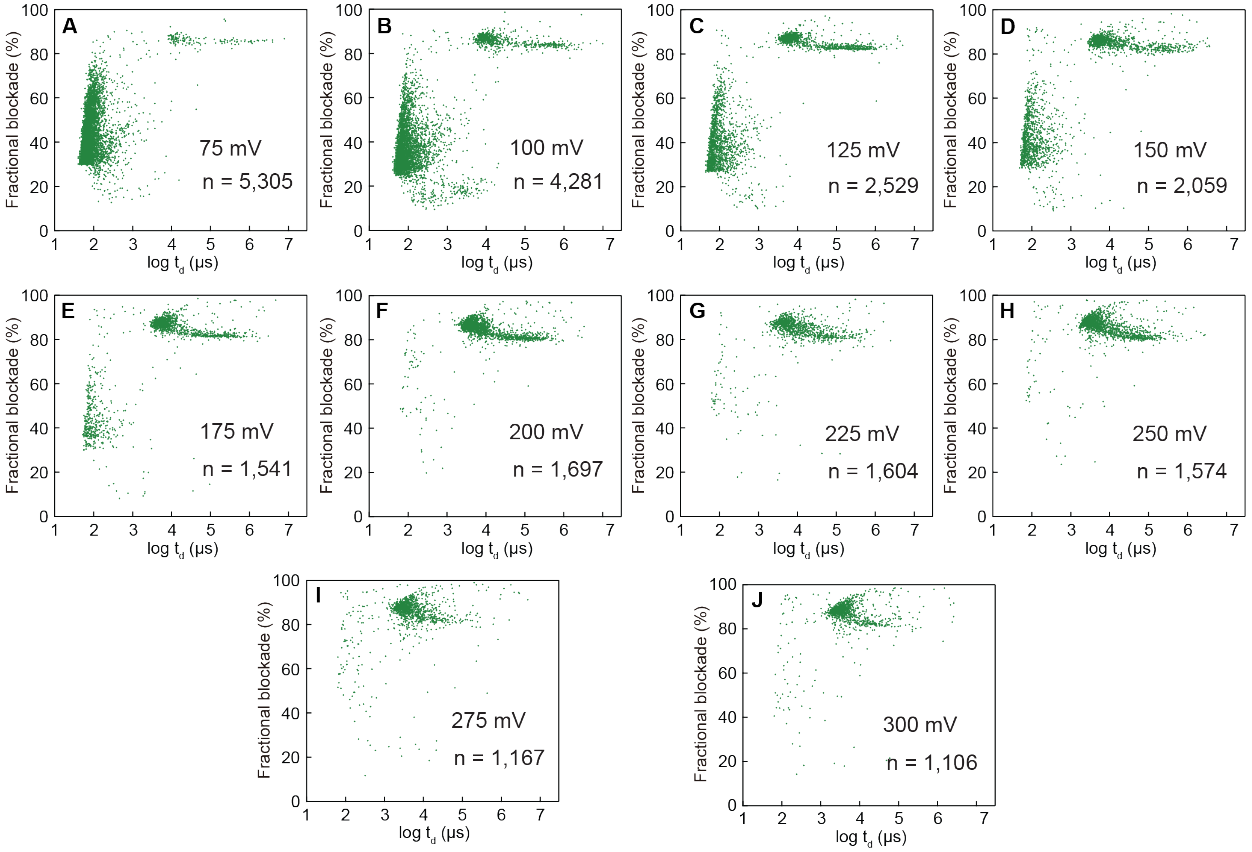


**Figure S7.** Fractional current blockade vs. dwell time scatters for MBP-D10 (0.35 μM) in 1.5 M GdmCl (1 M KCl, 10 mM Tris, pH 7.5), V = 75 – 300 mV.

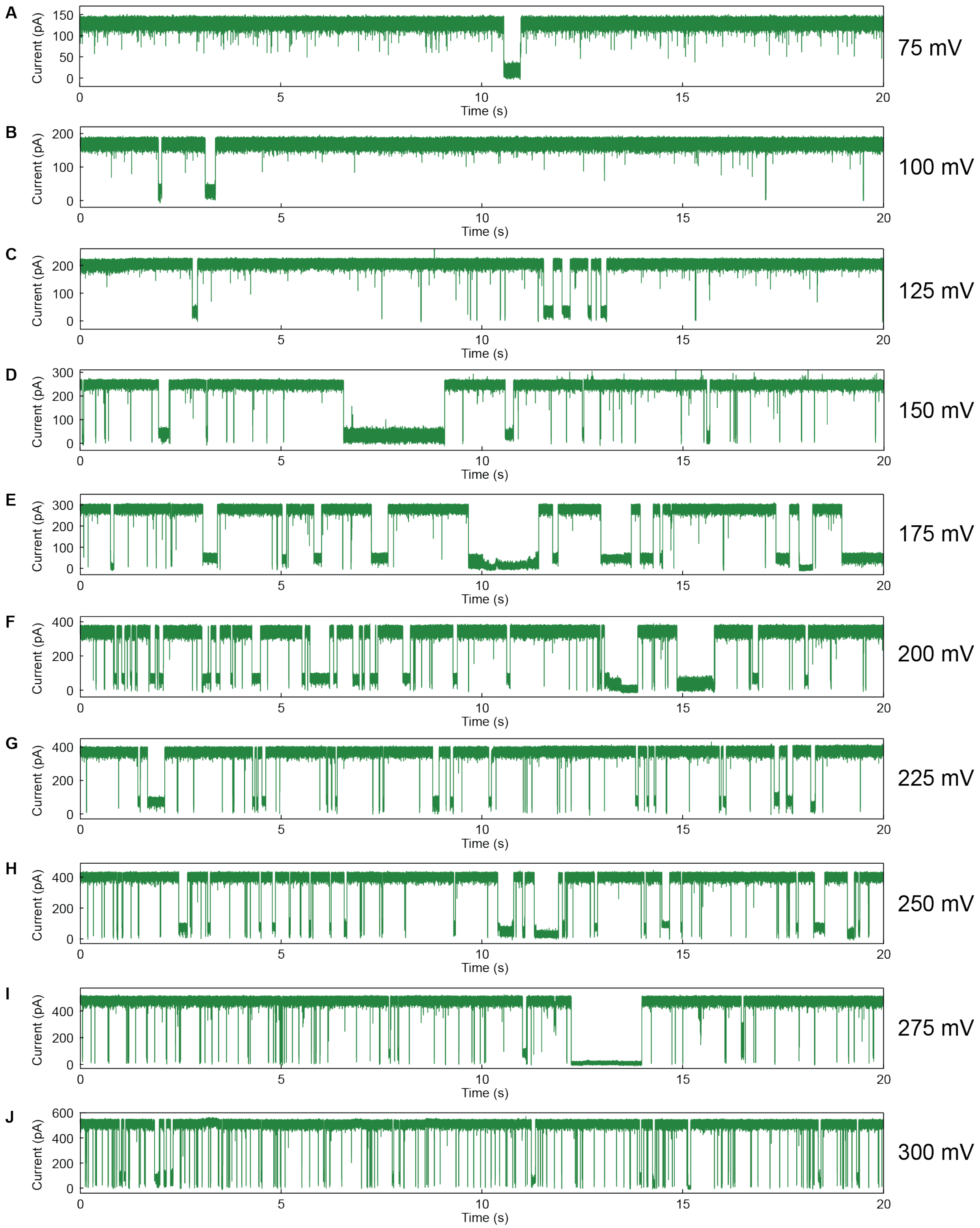


**Figure S8.** Current vs. time traces for MBP-D10 (0.35 μM) in 1.5 M GdmCl (1 M KCl, 10 mM Tris, pH 7.5), V = 75 – 300 mV.

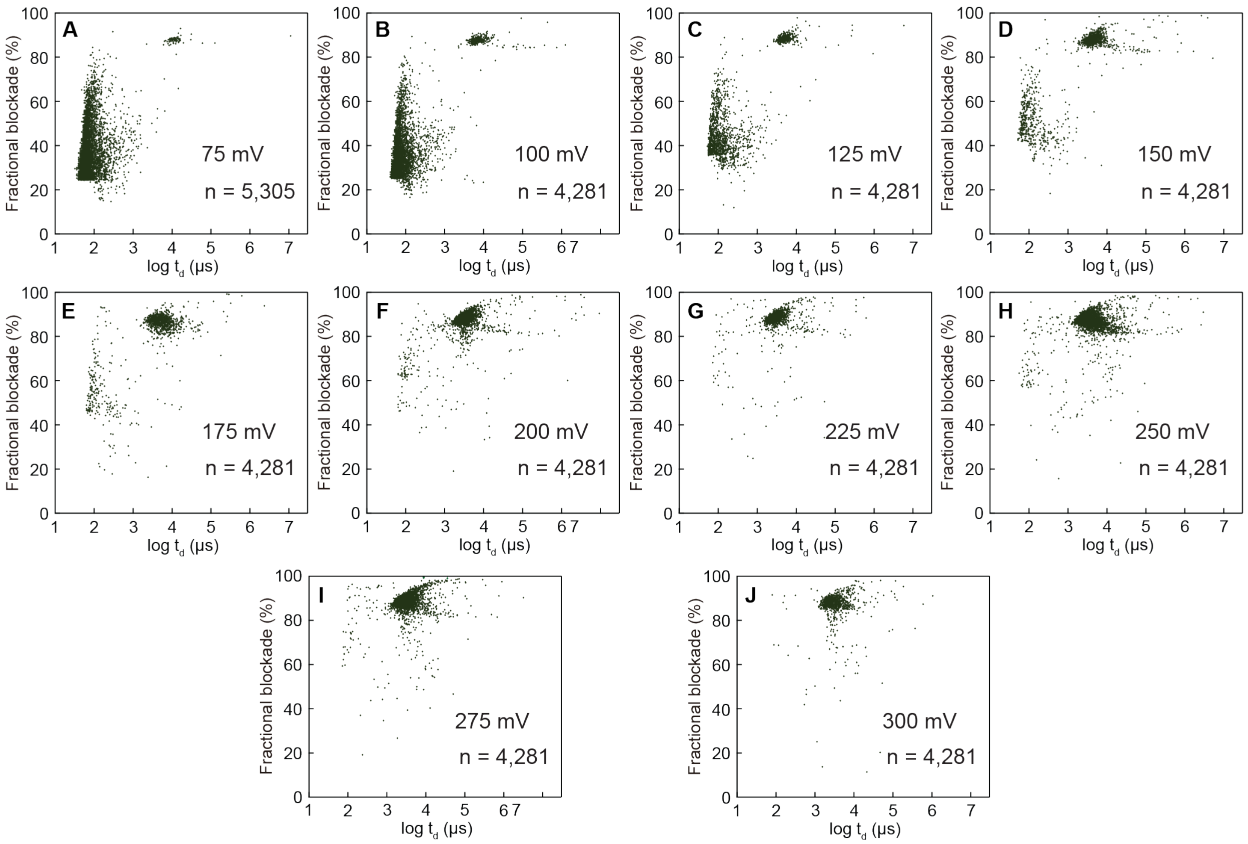


**Figure S9.** Fractional current blockade vs. dwell time scatters for MBP-D10 (0.35 μM) in 2.0 M GdmCl (1 M KCl, 10 mM Tris, pH 7.5), V = 75 – 300 mV.

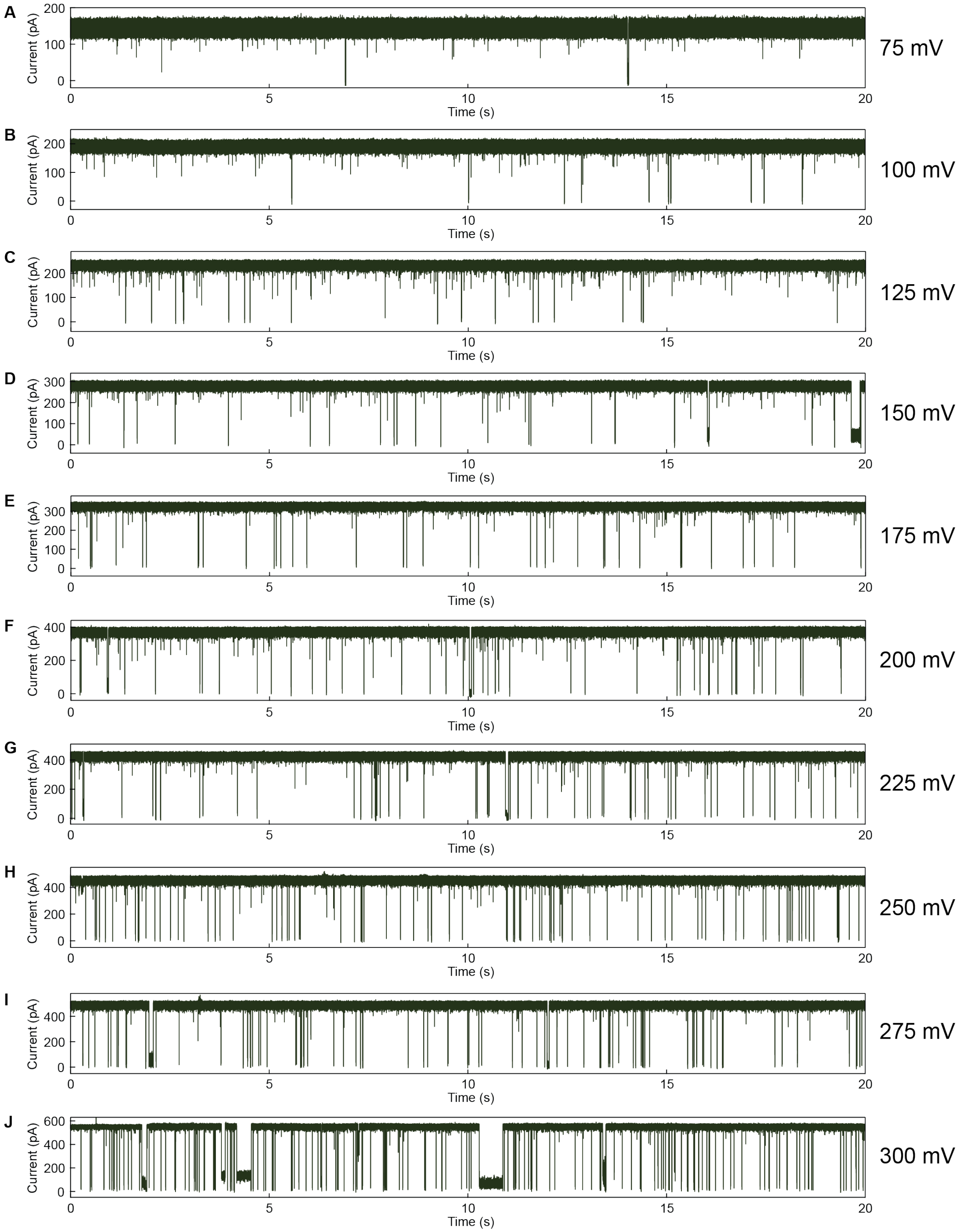


**Figure S10.** Current vs. time traces for MBP-D10 (0.35 μM) in 2.0 M GdmCl (1 M KCl, 10 mM Tris, pH 7.5), V = 75 – 300 mV.

#### Section 3: Extra data and analysis for diMBP-D10

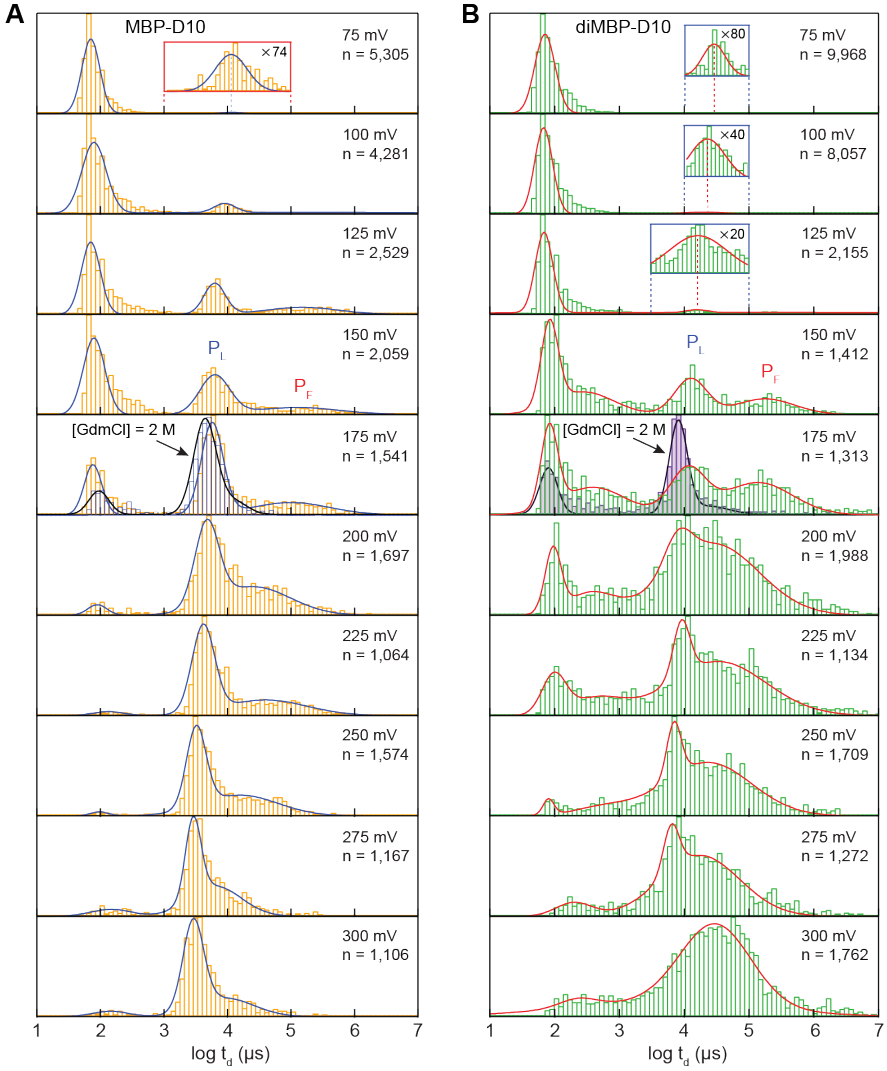


**Figure S11.** Dwell time histograms of **A)** MBP-D10 (0.35 μM) and **B)** diMBP-D10 (0.35 μM) at 75-300 mV and log-normal fits (1.5 M GdmCl, 1 M KCl). In the 75-175 mV voltage range, a significant fraction of events has ~100 μs dwell times, and as voltage increases further, a greater fraction of the events have longer dwell times in the range of 1-1000 ms, concentrated in two populations that correspond to $P_F$ and $P_L$. Strikingly, the position of the $P_L$ population shifts monotonically toward faster dwell times as voltage is increased for both monomeric and dimeric MBP (top to bottom). To highlight the impact of higher GdmCl concentrations on complete protein unfolding, we show dwell time distributions at 175 mV in 2 M GdmCl (1 M KCl) for both monomeric (n=1155) and dimeric (n=1151) MBP (blue and purple histograms, respectively).

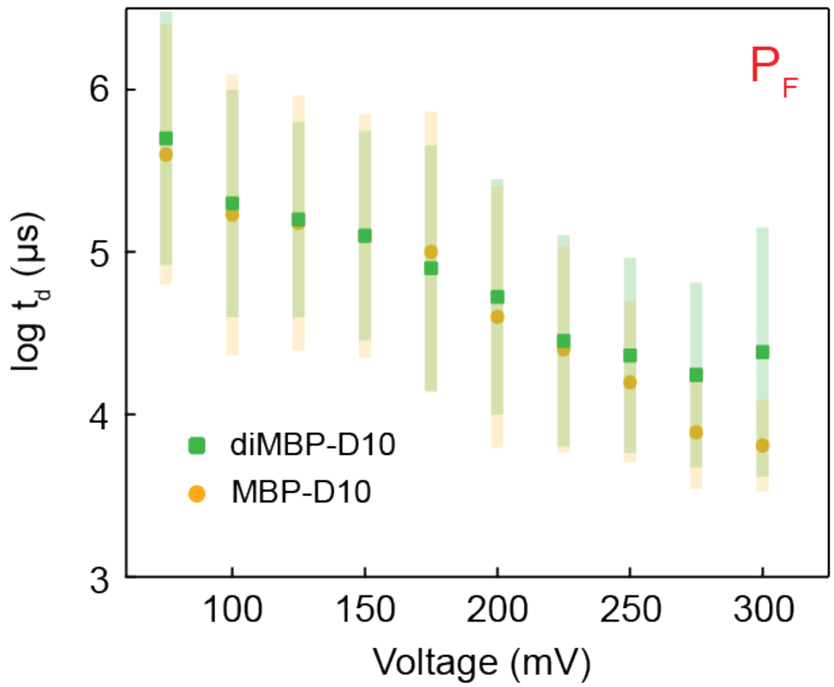


**Figure S12.** Mean dwell time vs. voltage for the $P_F$ populations of MBP-D10 and diMBP-D10, respectively (error bars represent the FWHM of the distribution fits). Buffer: 1 M KCl, 1.5 M GdmCl, 10 mM Tris, pH 7.5.

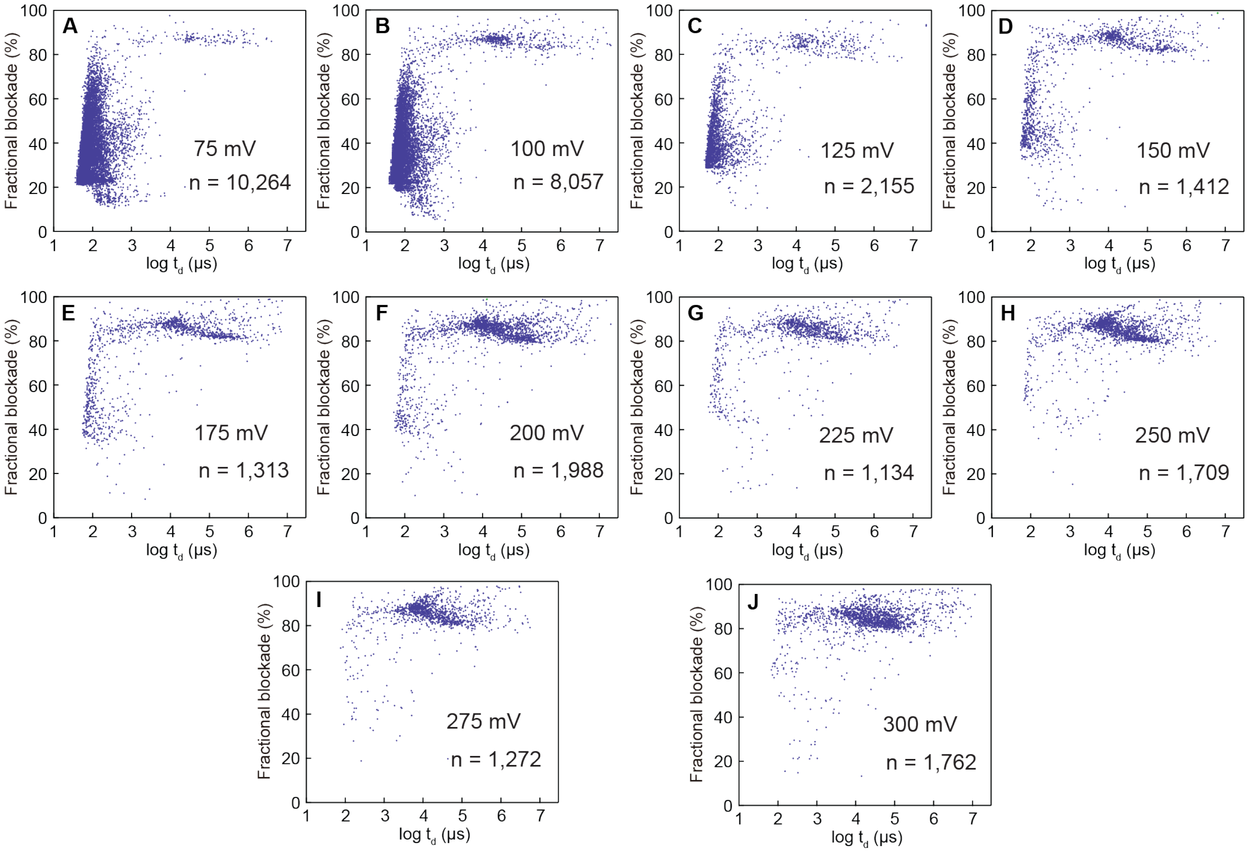


**Figure S13.** Fractional current blockade vs. dwell time scatters for diMBP-D10 (0.35 μM) in 1.5 M GdmCl (1 M KCl, 10 mM Tris, pH 7.5), V = 75 – 300 mV.

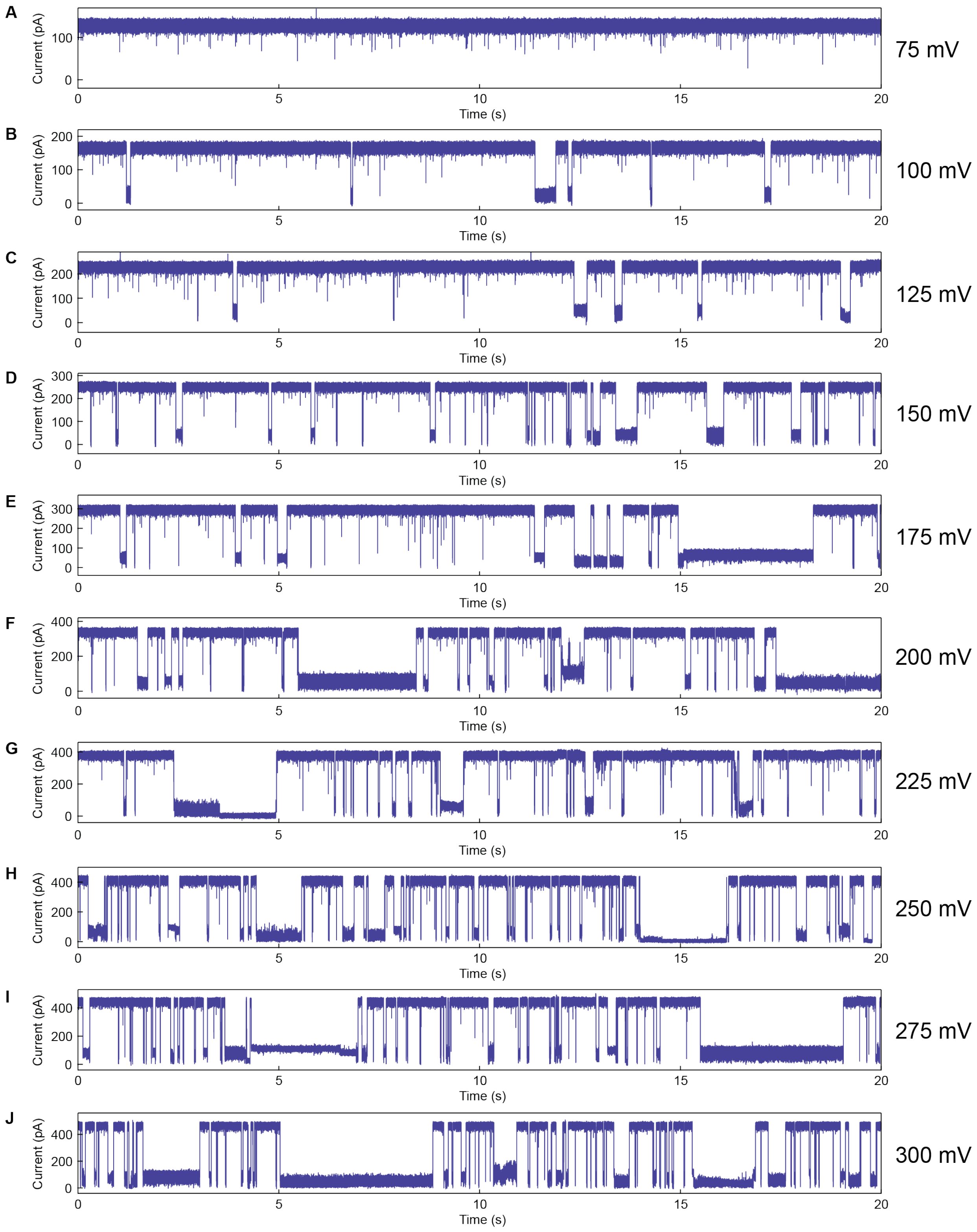


**Figure S14.** Current vs. time traces for diMBP-D10 (0.35 μM) in 1.5 M GdmCl (1 M KCl, 10 mM Tris, pH 7.5), V = 75 – 300 mV.

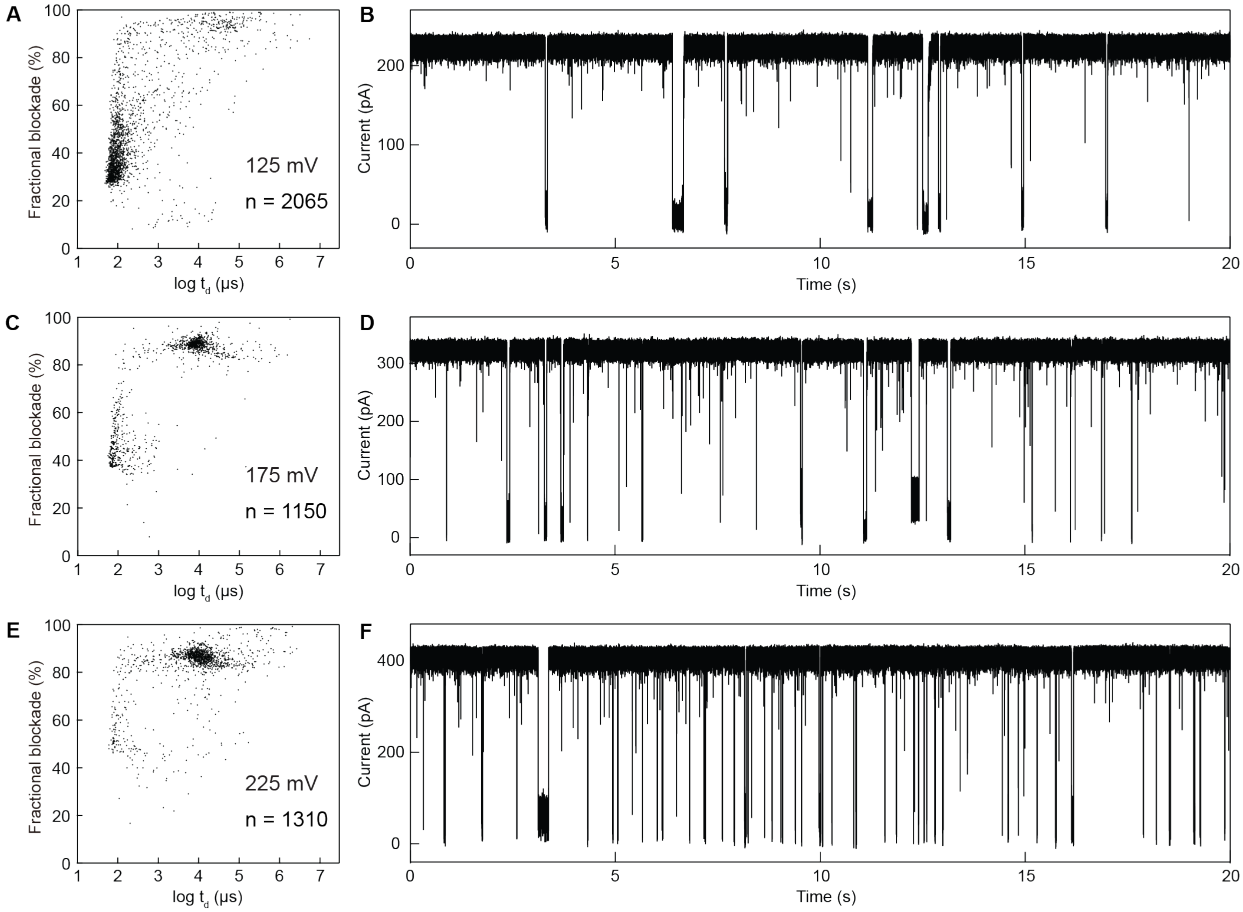


**Figure S15.** Fractional current blockade vs. dwell time scatters (left) and current vs. time traces (right) for diMBP-D10 (0.35 μM) in 2.0 M GdmCl (1 M KCl, 10 mM Tris, pH 7.5), V = 125, 175, 225 mV.

#### Section 4: Other experimental data

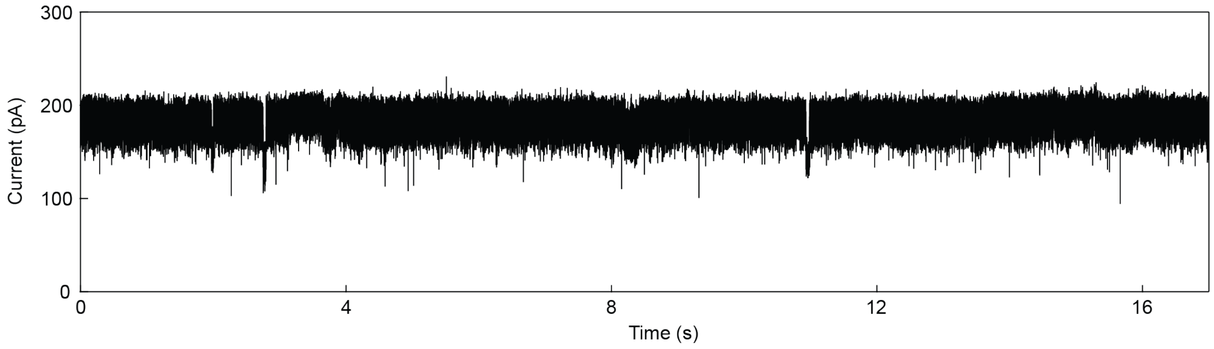


**Figure S16.** Current vs. time trace for for MBP-D10 (0.35 μM) without GdmCl in KCl, 10mM Tris, pH 7.5, V = 175 mV.

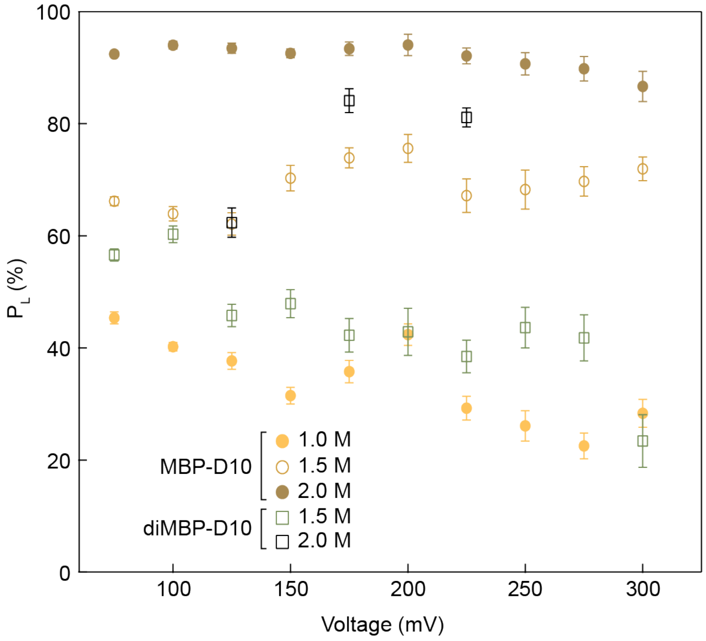


**Figure S17.** $P_L$ percentage vs. voltage for MBP-D10 (0.35 μM) in 1.0, 1.5, 2.0 M GdmCl and diMBP-D10 (0.35 μM) in 1.5, 2.0 M GdmCl (all buffers include 1 M KCl, 10 mM Tris, pH 7.5).

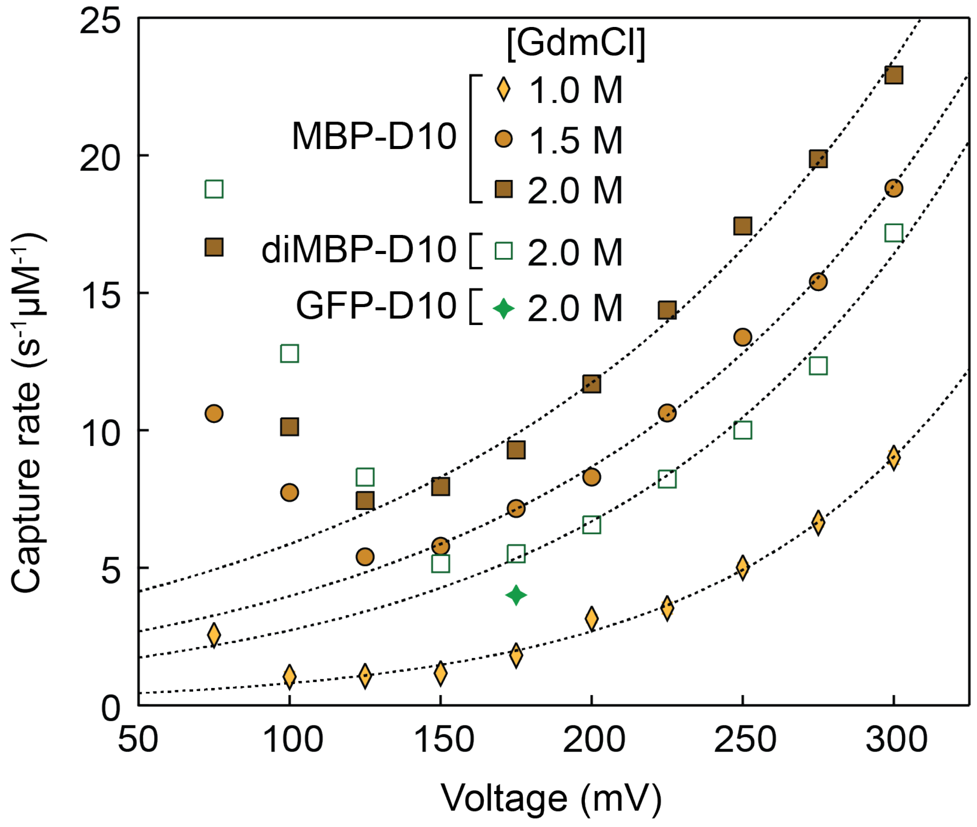


**Figure S18.** Concentration-normalized capture rates as a function of voltage for MBP-D10 (in 1.0 M, 1.5 M and 2.0 M GdmCl + 1 M KCl) and diMBP-D10 (in 1.5 M GdmCl + 1 M KCl). The green star mark shows capture rate for GFP-D10 at 175 mV (in 2.0 M GdmCl + 1 M KCl). Error bars are smaller than the marker sizes in all cases. Dashed lines are exponential fits to the data in the range shown, where low voltage data was excluded (since the high capture rates mainly come from collisions between analytes and α-hemolysin vestibule).

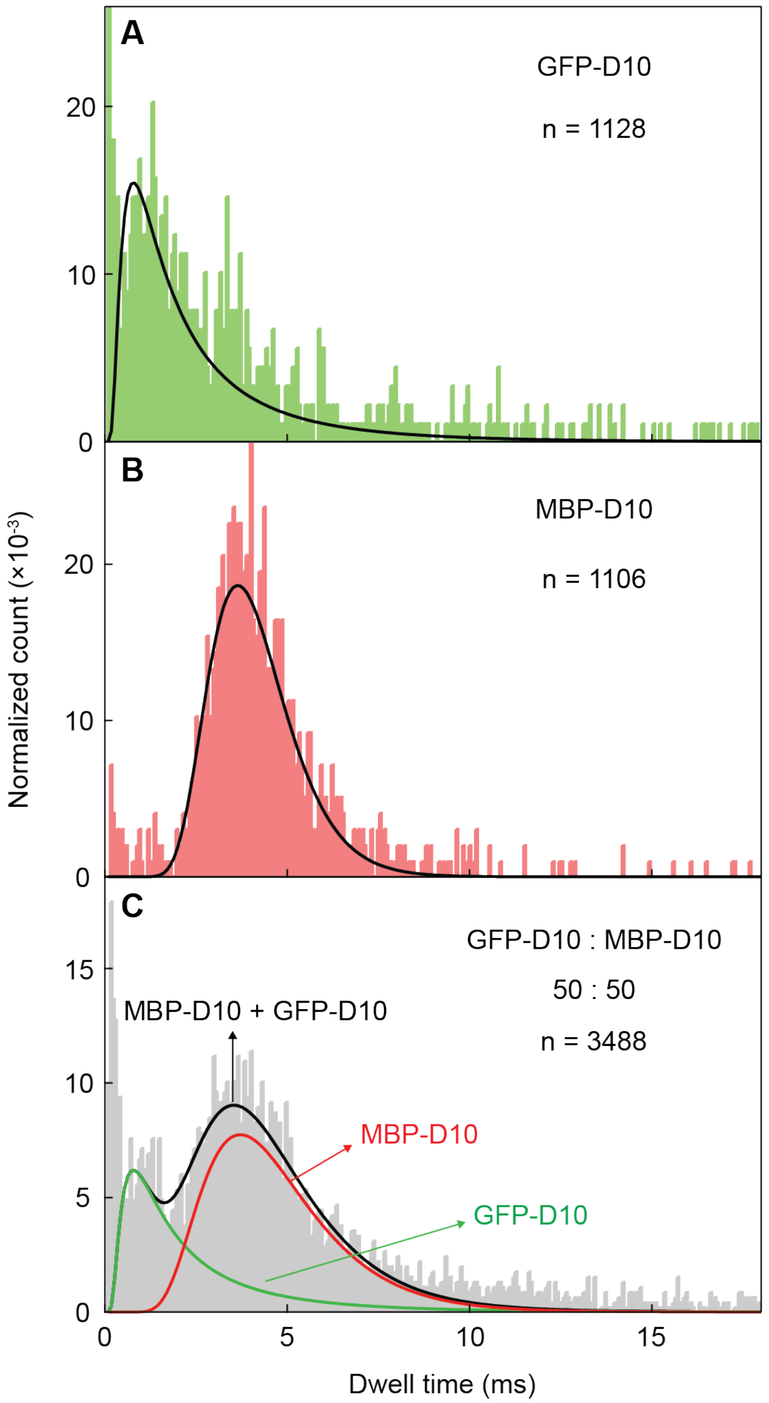


**Figure S19.** Dwell time histograms (normalized by event numbers) in 2 M GdmCl (1 M KCl, 10 mM Tris, pH 7.5) for **A)** 0.35 μM MBP-D10, **B)** 0.35 μM GFP-D10 and **C)** mixture of 0.35 μM MBP-D10 and 0.35 μM GFP-D10 with fitting lines based on 1D drift-diffusion model (details in **Supplementary Note 2** and **Table S7, Exp. 2**).

**Supplementary Note 2: Estimate of the event ratio in the MBP-D10/GFP-D10 mixture**

Only considering the translocation events with dwell time range $0.3 < t_d < 20$ ms (not considering the collisions and extremely long events), the dwell time histograms in **Figure S19, panel A and B** are fitted with 1D drift-diffusion function

$$P(t) = \frac{h_{eff}}{\sqrt{4\pi D t^3}} exp\left(-\frac{(h_{eff} - vt)^2}{4Dt}\right) \quad \textbf{(Eq. S1)}$$

that has been used in previous DNA and protein translocation studies[2-5]. $h_{eff}$ is the effective height of the nanopore, $D$ is the molecule diffusion constant, and $v$ is the protein drift velocity. **Figure S19C** shows the dwell time histogram for the mixture of 0.35 μM GFP-D10 and 0.35 μM MBP-D10. Since GFP-D10 and MBP-D10 events are independent of each other, we can fit the histogram using formula

$$P(t) = \frac{h_{eff}}{\sqrt{4\pi D t^3}} exp\left(-\frac{(h_{eff} - vt)^2}{4Dt}\right) + \frac{h'_{eff}}{\sqrt{4\pi D' t^3}} exp\left(-\frac{(h'_{eff} - v't)^2}{4D't}\right) \quad \textbf{(Eq. S2)}$$

where coefficients $h_{eff}$, $D$, $v$ are for GFP-D10 and $h'_{eff}$, $D'$, $v'$ are for MBP-D10. Individual GFP-D10 (green) and MBP-D10 (red) fitting lines are shown based on the same coefficients in **Equation. S2**, with the ratio of areas (calculated based on integrals of the fitting lines) between GFP-D10 and MBP-D10 being 32.6:67.4, which is consistent with the result in **Figure 5D**.

### Section 5: MD simulations

**Table S3.** Fragmentation of the maltose-binding protein (MBP) into seven peptides for MD simulations. Negatively charged residues are highlighted in blue and positively charged residues are shown in red. The amino acid sequence of MBP was taken from the Protein Data Bank (PDB ID: 1JW4).

| Fragment | Residue ID | Sequence | Charge (e) |
|---|---|---|---|
| 1 | 1 to 53 | KIEEGKLVIWINGDKGYNGLAEVGKKFEKDTGIKVTVEHPDKLEEKFPQVAAT | -1 |
| 2 | 54 to 106 | GDGPDIIFWAHDRFGGYAQSGLLAEITPDKAFQDKLYPFTWDAVRYNGKLIAY | -2 |
| 3 | 108 to 159 | PIAVEALSLIYNKDLLPNPPKTWEEIPALDKELKAKGKSALMFNLQEPYFTWP | -1 |
| 4 | 160 to 212 | LIAADGGYAFKYENGKYDIKDVGVDNAGAKAGLTFLVDLIKNKHMNADTDYSI | -2 |
| 5 | 213 to 265 | AEAAFNKGETAMTINGPWAWSNIDTSKVNYGVTVLPTFKGQPSKPFVGVLSAG | 1 |
| 6 | 266 to 318 | INAASPNKELAKEFLENYLLTDEGLEAVNKDKPLGAVALKSYEEELAKDPRIA | -4 |
| 7 | 319 to 370 | ATMENAQKGEIMPNIPQMSAFWYAVRTAVINAASGRQTVDEALKDAQTRITK | 1 |

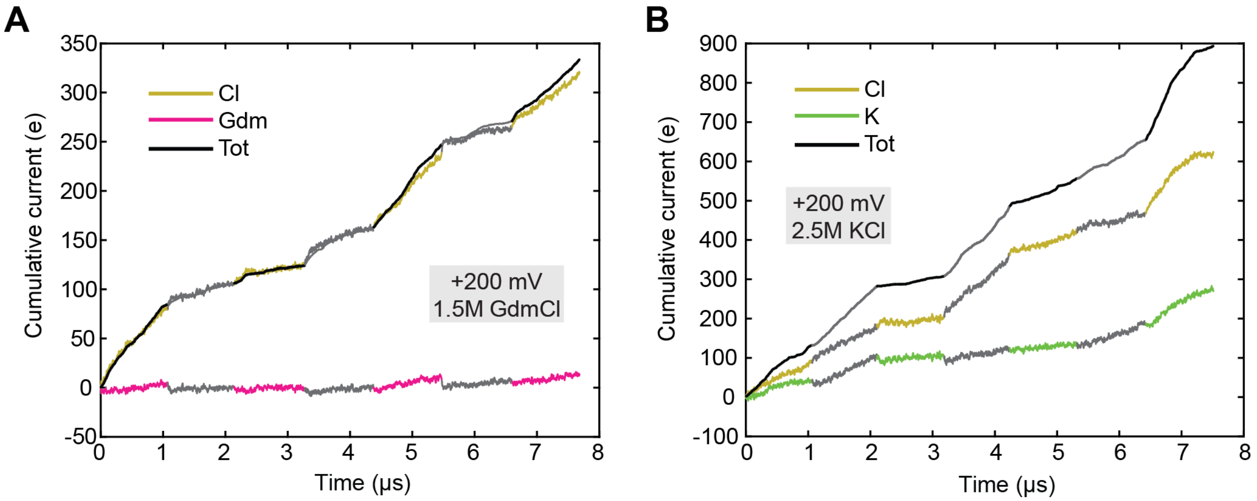


**Figure S20.** MD simulation of α-hemolysin systems containing fragments of MBP. Total charge carried by the ion species through the α-hemolysin nanopore is plotted versus simulation time for seven independent MD simulations carried out under a +200 mV bias for 1.5 M GdmCl **(A)** and 2.5 M KCl electrolyte **(B)** conditions. Each trace is shown using two alternating colors to indicate data from the seven MD trajectories differing by the sequence and conformation of the MBP fragment. The traces were added consecutively to appear as a continuous permeation trace.

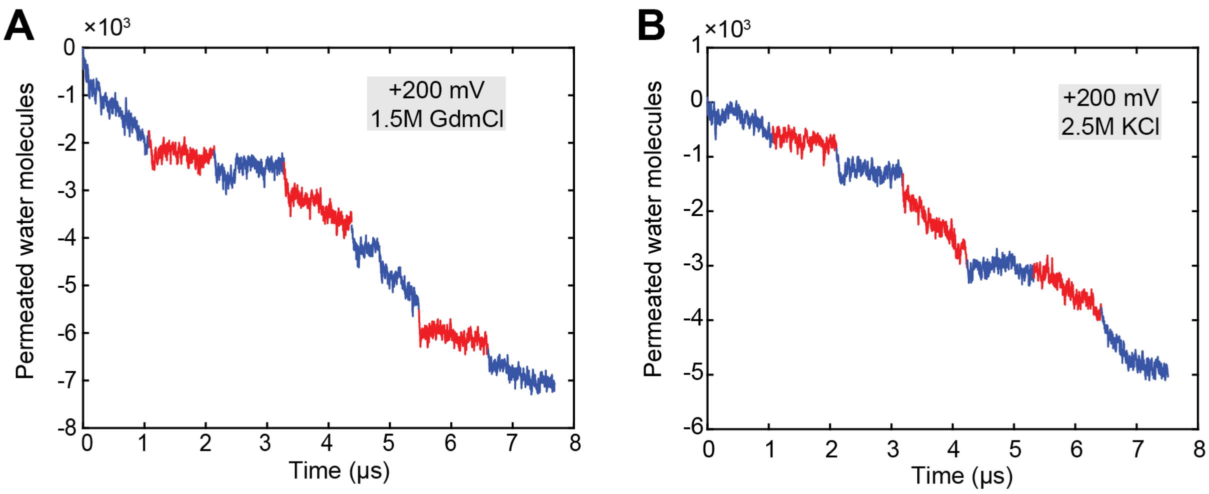


**Figure S21.** Simulated electro-osmotic flow in α-hemolysin systems containing fragments of MBP. The number water molecules permeated through the α-hemolysin constriction (residues 111, 113, and 147) is plotted as a function of simulation time for MD simulations carried out under a +200 mV bias for the 1.5 M GdmCl **(A)** and 2.5 M KCl **(B)** electrolyte conditions. Negative values indicate transport in the direction opposite that of the *z*-axis (defined in **Figure 3A**). Results for the seven independent simulations are shown using alternating colors. The traces were added consecutively to appear as a continuous permeation trace.

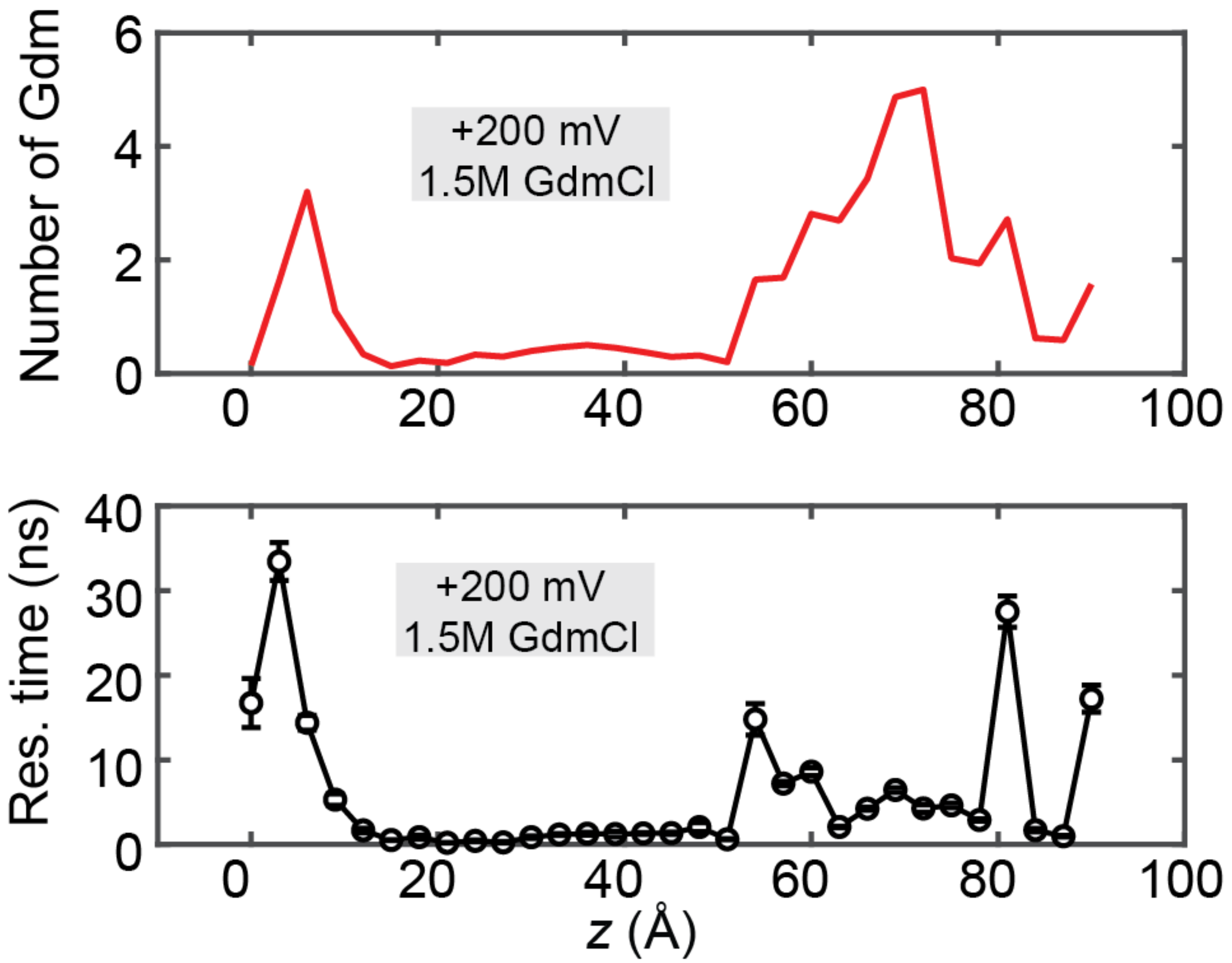


**Figure S22.** Gdm binding to the inner surface of α-hemolysin in MD simulations of the α-hemolysin systems containing MBP fragments. The top row shows the number of Gdm ions located within 3 Å of the nanopore inner surface. The bottom row shows the average residence time of the Gdm ions near the α-hemolysin nanopore surface. The *z*-axis is defined in **Figure 3A**. The data were averaged over the seven replica simulations at the same electrolyte condition.

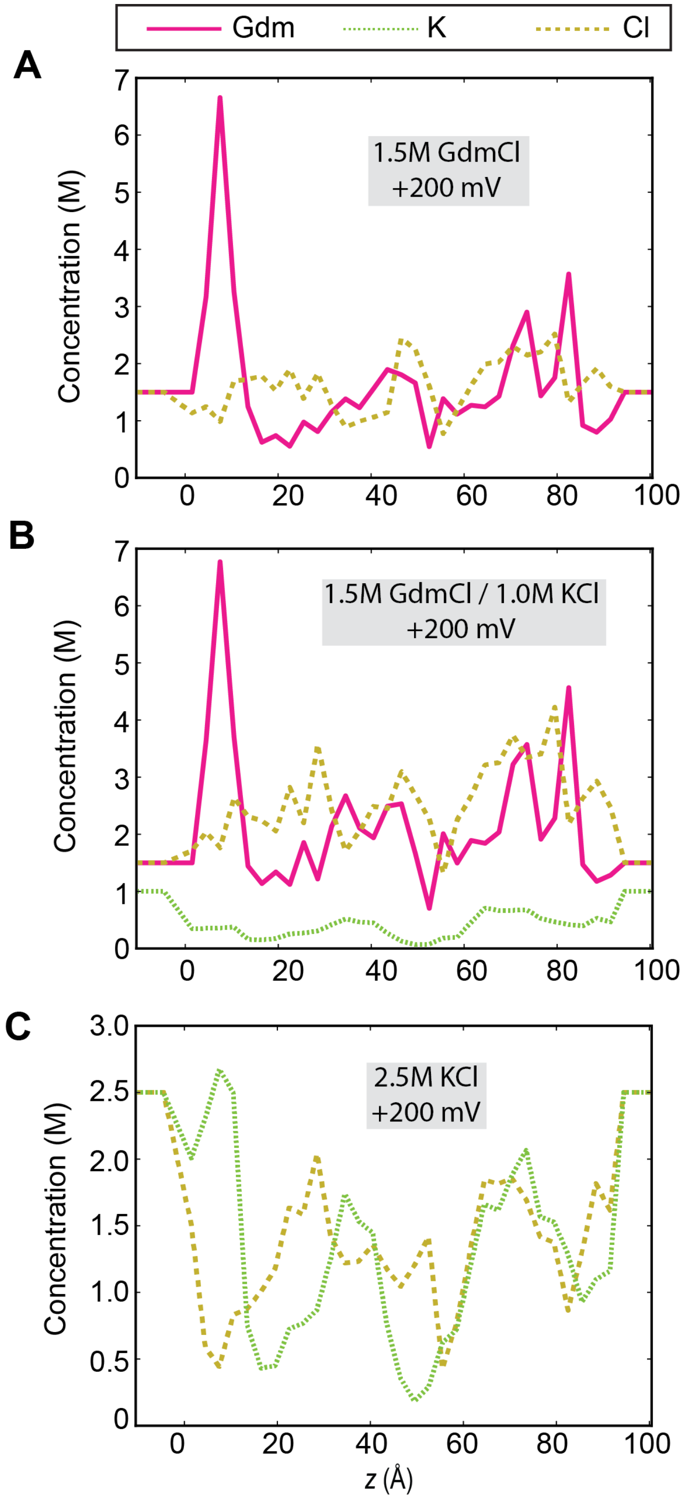


**Figure S23.** Local concentration of ions along the transmembrane nanopore observed in MD simulations of the α-hemolysin systems containing fragments of MBP. For each electrolyte condition, the concentrations were averaged the seven replica simulations in 3 Å bin along the nanopore axis (the *z* axis). The electrolyte and applied bias conditions are specified in each panel.

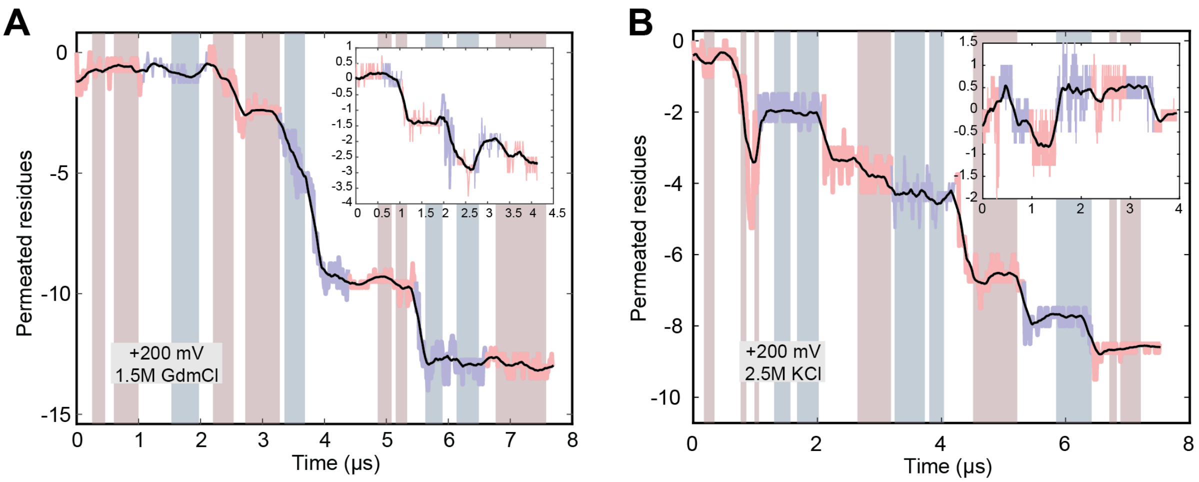


**Figure S24.** Simulated transport of MBP fragments. The number of amino acid residues permeated through the α-hemolysin constriction (residues 111, 113, and 147) is plotted as a function of simulations for seven independent MD simulations carried out under a +200 mV bias for the 1.5 M GdmCl **(A)** and 2.5 M KCl **(B)** electrolyte conditions. Negative values indicate transport in the direction opposite that of the *z*-axis (defined in **Figure 3A**). Results for the seven independent simulations are shown using alternating colors. The traces were added consecutively to appear as a continuous permeation trace. Highlights indicate the parts of the trajectories where the peptide density within the stem (8<z<45 Å) of α-hemolysin is constant; the inset shows consecutive addition of the highlighted regions.

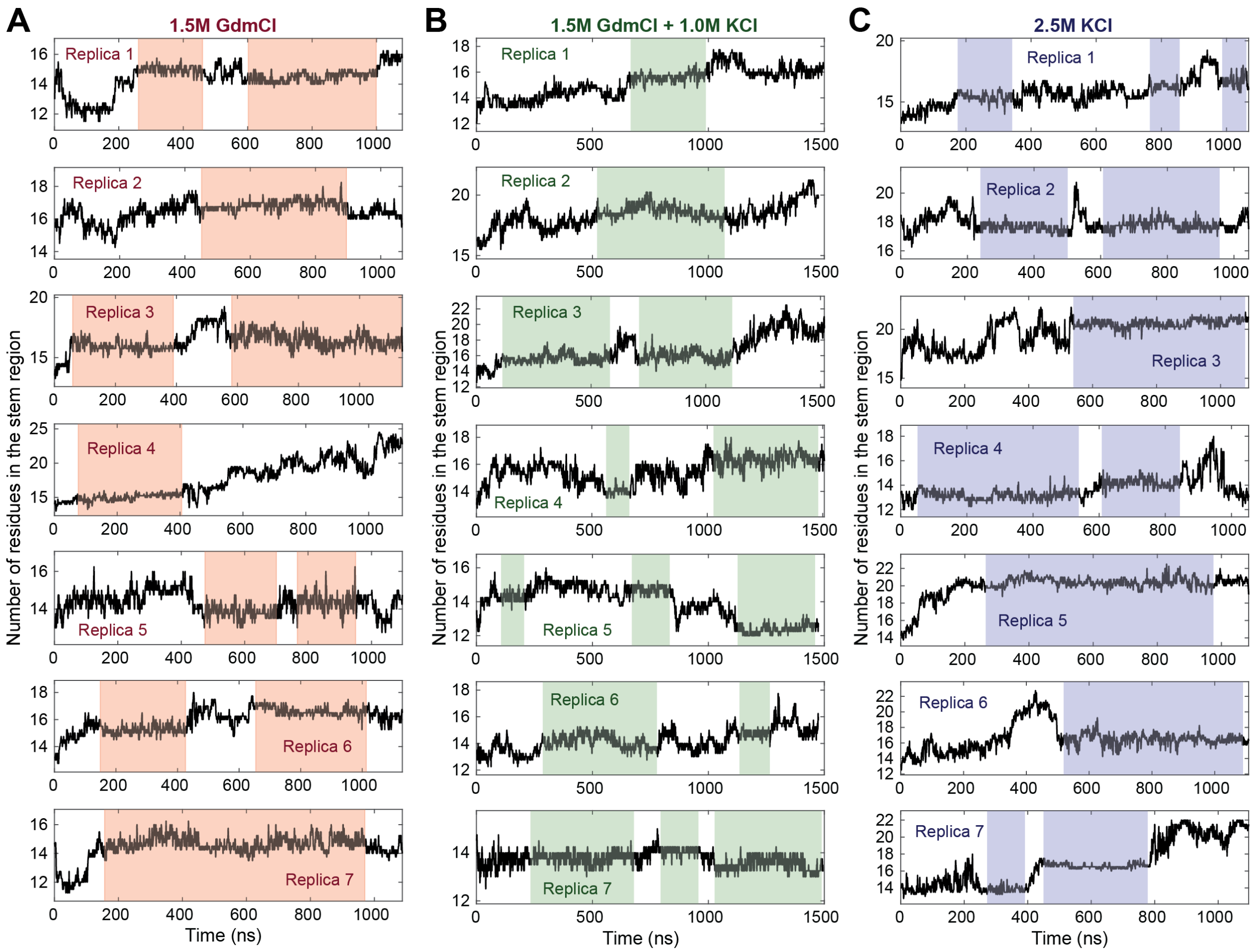


**Figure S25.** Number of residues confined within the stem region of α-hemolysin nanopore. Panels A, B and C correspond to simulations carried out at 1.5 M GdmCl **(A)**, 1.5 M GdmCl / 1.0 M KCl **(B)** and 2.5 M KCl **(C)** electrolyte solutions. Each plots shows data for one independent simulation. The highlights indicate the parts of the trajectories where the peptide density is nearly constant. These regions were used to calculate the peptide translocation rate displayed in **Figure 3H**.

### Section 6: Machine learning analysis

**Table S4.** Selection and processing parameters for translocation events used in the directionality analysis via Soft-DTW (shown in **Figure 4A**).

| MBP Variant | # Total events | Event Selection Criteria | | | # Passing events | Calculated smoothing factor γ |
|---|---|---|---|---|---|---|
| | | Minimum duration (ms) | Maximum duration (ms) | Maximum current (pA) | | |
| MBP-D10 | 854 | 3.0 | 5.0 | 130 | 331 | 1.642 |
| D10-MBP | 488 | 3.0 | 5.0 | 130 | 174 | 2.351 |
| diMBP-D10 | 713 | 6.0 | 9.0 | 130 | 288 | 2.749 |

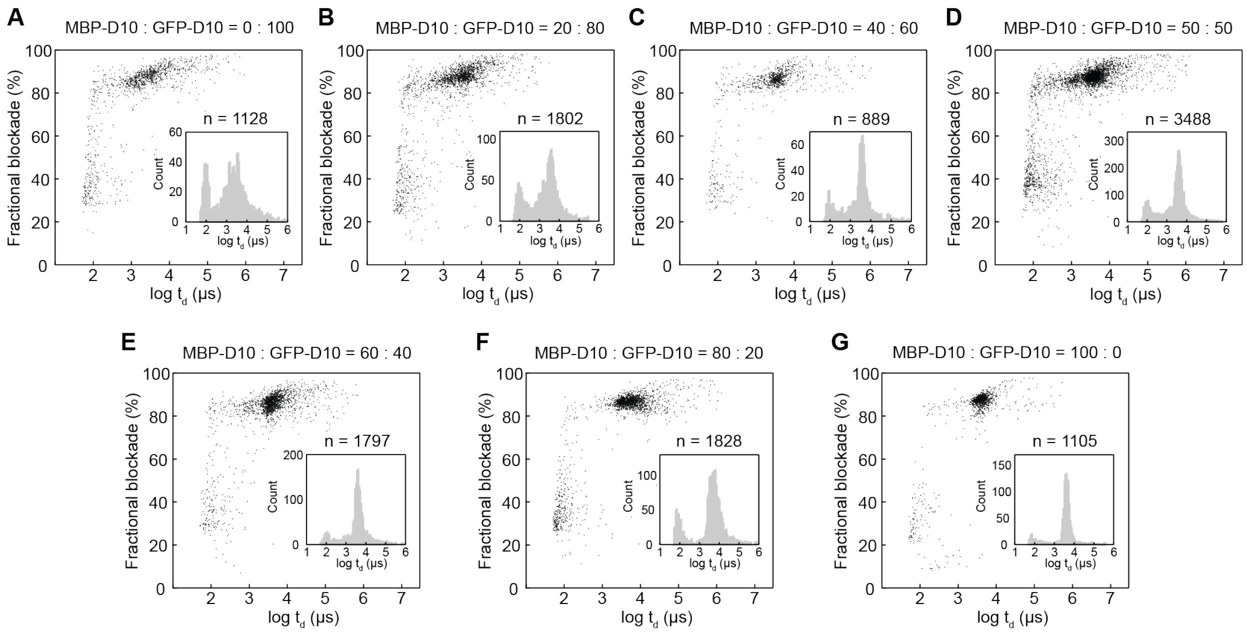


**Figure S26.** Fractional current blockade vs. dwell time scatter plots and corresponding dwell time histograms for different GFP-D10:MBP-D10 concentration ratios. All experiments were performed in 1.0 M KCl, 2.0 M GdmCl, 10 mM Tris, pH 7.5 and under a 175 mV bias applied to the *trans* chamber.

**Table S5.** Sample size of translocation events parsed for each protein type from pure experiments with dwell times that met the conditions of being 1) longer than 0.3 ms and 2) shorter than 20 ms for training and testing GBC models.

| Protein Type | Minimum Duration (ms) | Maximum Duration (ms) | Total number of events |
|---|---|---|---|
| MBP-D10 | 0.3 | 20 | 1760 |
| D10-MBP | 0.3 | 20 | 488 |
| diMBP-D10 | 0.3 | 20 | 713 |
| GFP-D10 | 0.3 | 20 | 979 |

**Table S6.** Sample size of translocation events parsed from MBP-D10:GFP-D10 mixture experiments with dwell times that met the conditions of being 1) longer than 0.3 ms and 2) shorter than 20 ms. The trained GBC models generated for discrimination of MBP-D10 and GFP-D10 were applied and used to classify these events with an associated probability score.

| Mixture Ratio (MBP-D10:GFP-D10) | Minimum Duration (ms) | Maximum Duration (ms) | Total number of events |
|---|---|---|---|
| 20:80 | 0.3 | 20 | 1221 |
| 40:60 | 0.3 | 20 | 640 |
| 50:50 (Exp 1) | 0.3 | 20 | 1160 |
| 50:50 (Exp 2) | 0.3 | 20 | 2491 |
| 50:50 (Combined) | 0.3 | 20 | 3651 |
| 60:40 | 0.3 | 20 | 1387 |
| 80:20 | 0.3 | 20 | 1295 |

**Table S7.** Predicted ratios by trained GBC model on unlabeled events parsed from MBP-D10:GFP-D10 mixture experiments (sample size for each experiment is shown in **Table S6**). The ratio results are the mean and standard deviation of 9 GBC models that were trained and tested on a reshuffled dataset comprised of labeled MBP-D10 and GFP-D10 samples from pure experiments. The ratio results shown in this table are presented in **Figure 5D**. The mean and standard deviation of the probability score for all events classified as MBP-D10 or GFP-D10 by a trained GBC model is shown for each experiment.

| Mixture Ratio (MBP-D10:GFP-D10) | MBP-D10 Ratio (%) | GFP-D10 Ratio (%) | Total MBP-D10 Probability Score (%) | Total GFP-D10 Probability Score (%) |
|---|---|---|---|---|
| 20:80 | 43.7 ± 0.6 | 56.3 ± 0.6 | 92.2 ± 12 | 92.1 ± 13 |
| 40:60 | 55.4 ± 1.0 | 44.6 ± 1.0 | 91.9 ± 12 | 91.5 ± 12 |
| 50:50 (Exp 1) | 50.7 ± 0.6 | 49.3 ± 0.6 | 89.8 ± 13 | 91.8 ± 13 |
| 50:50 (Exp 2) | 67.4 ± 0.4 | 32.6 ± 0.4 | 94.7 ± 10 | 91.1 ± 13 |
| 50:50 (combined) | 61.8 ± 0.5 | 38.2 ± 0.5 | 93.4 ± 11 | 91.1 ± 13 |
| 60:40 | 67.8 ± 1.3 | 32.2 ± 1.3 | 92.2 ± 12 | 88.5 ± 15 |
| 80:20 | 86.2 ± 0.3 | 13.8 ± 0.3 | 94.9 ± 10 | 86.5 ± 16 |

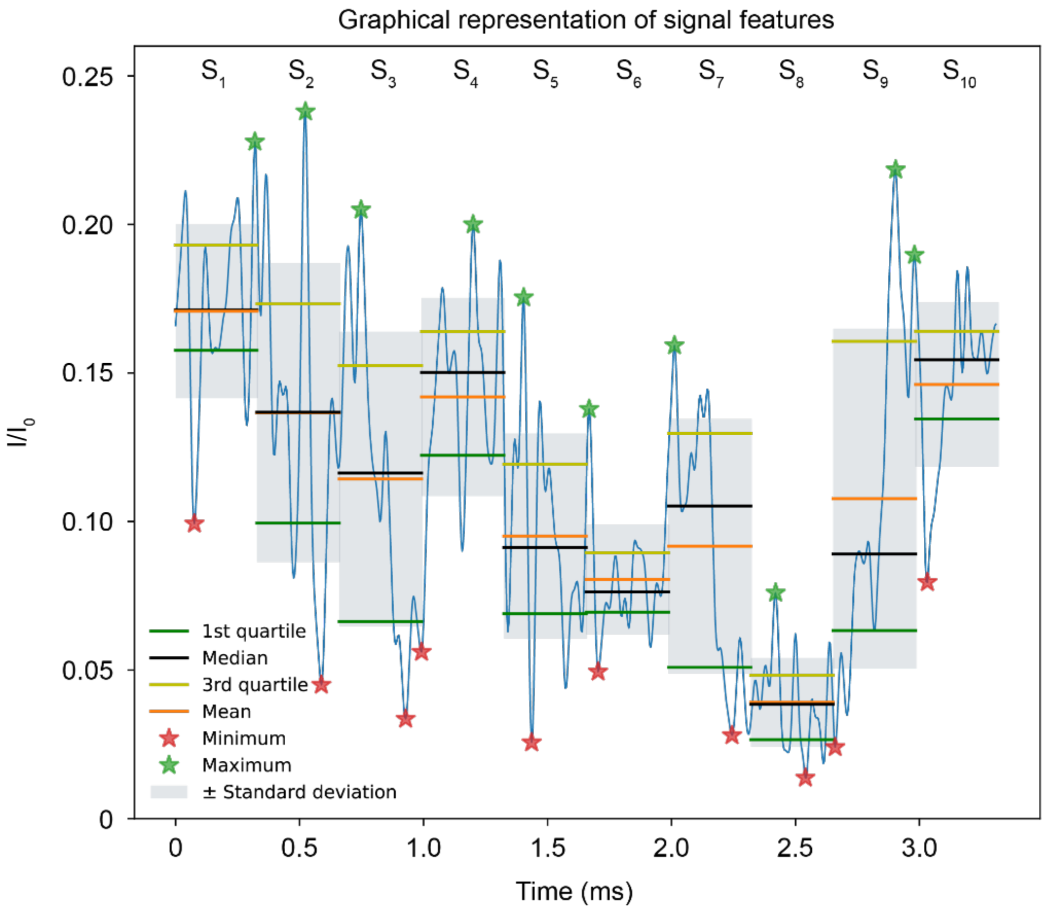


**Figure S27.** Graphical representation of the 7 statistical parameters (refer to legend) extracted from each segment (S1 to S10) of a full MBP-D10 translocation event (blue signal). Every translocation event used to train, test, and validate GBC models was divided into 10 segments of equal size, providing a total of 70 statistical features from every event. Note that every parameter was divided by the baseline current, $I_0$, as a normalization step to remove experiment-to-experiment variability during model fitting.

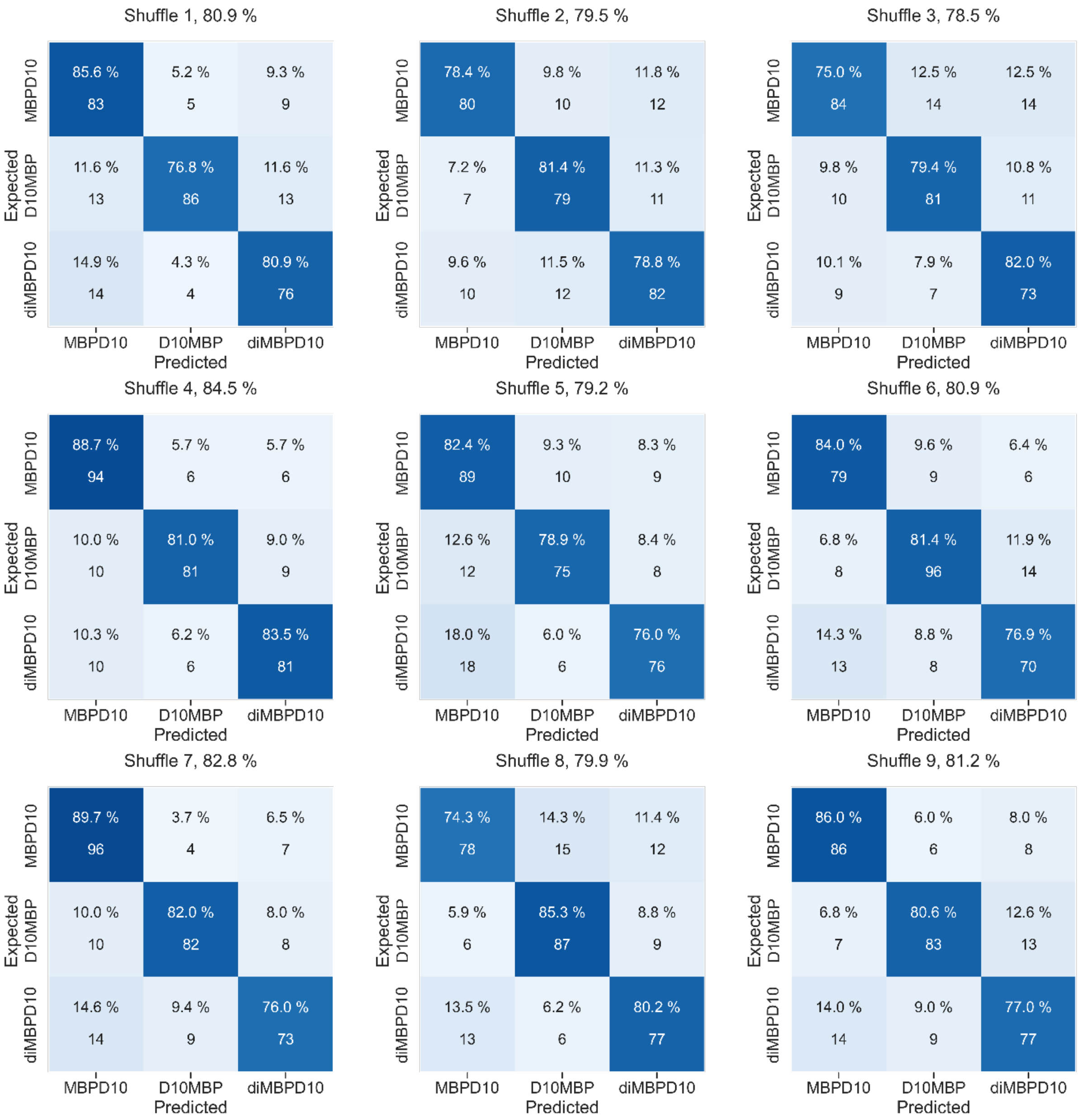


**Figure S28. MBP-D10 versus D10-MBP versus diMBP-D10 all iterations of training and testing.** Shown are the 9 confusion matrices, where each one is the result for a single training and testing iteration of a GBC model built for three-way discrimination of MBP variants. For each iteration, samples are randomly binned into the training or testing set, producing a unique combination of data for model fitting and evaluation. The distribution of model calls for each MBP variant in the test set are shown as percentages in each box, where each row sums to 100%. The number of samples (left column labels) called as a particular protein variant (bottom row labels) is shown below the percentage value. The results shown in the diagonal represents the correct classification. Displayed next to the shuffle number positioned above each confusion matrix is the overall classification of that GBC model iteration. The mean and standard deviation of the percent calls for all 9 GBC iterations are shown in **Figure 4B**.

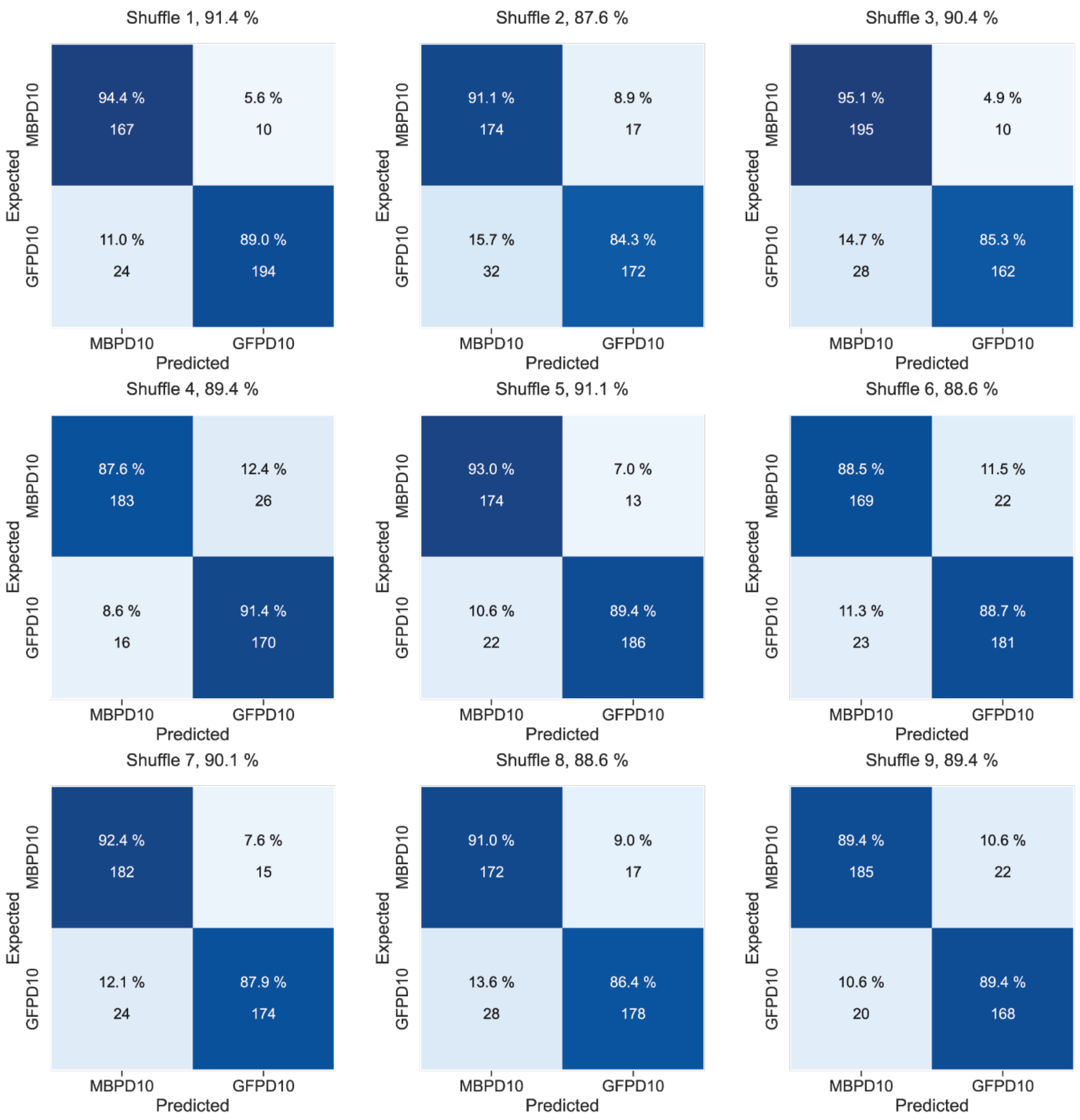


**Figure S29. MBP-D10 versus GFP-D10 all iterations of training and testing.** Shown are the 9 confusion matrices, where each one is the result of a single and unique training and testing iteration for a GBC model built for two-way discrimination of different protein classes (MBP-D10 and GFP-D10). For each iteration, samples are randomly binned into the training or testing set, producing a unique combination of data for model fitting and evaluation. The distribution of model calls for each protein class in the test set are shown as percentages in each box, where each row sums to 100%. The number of samples (left column labels) called as a particular protein class (bottom row labels) is shown below the percentage value. The results shown in the diagonal represents the correct classification. Displayed next to the shuffle number positioned above each confusion matrix is the overall classification of that GBC model iteration. The mean and standard deviation of the percent calls for all 9 GBC iterations are shown in **Figure 5E**.

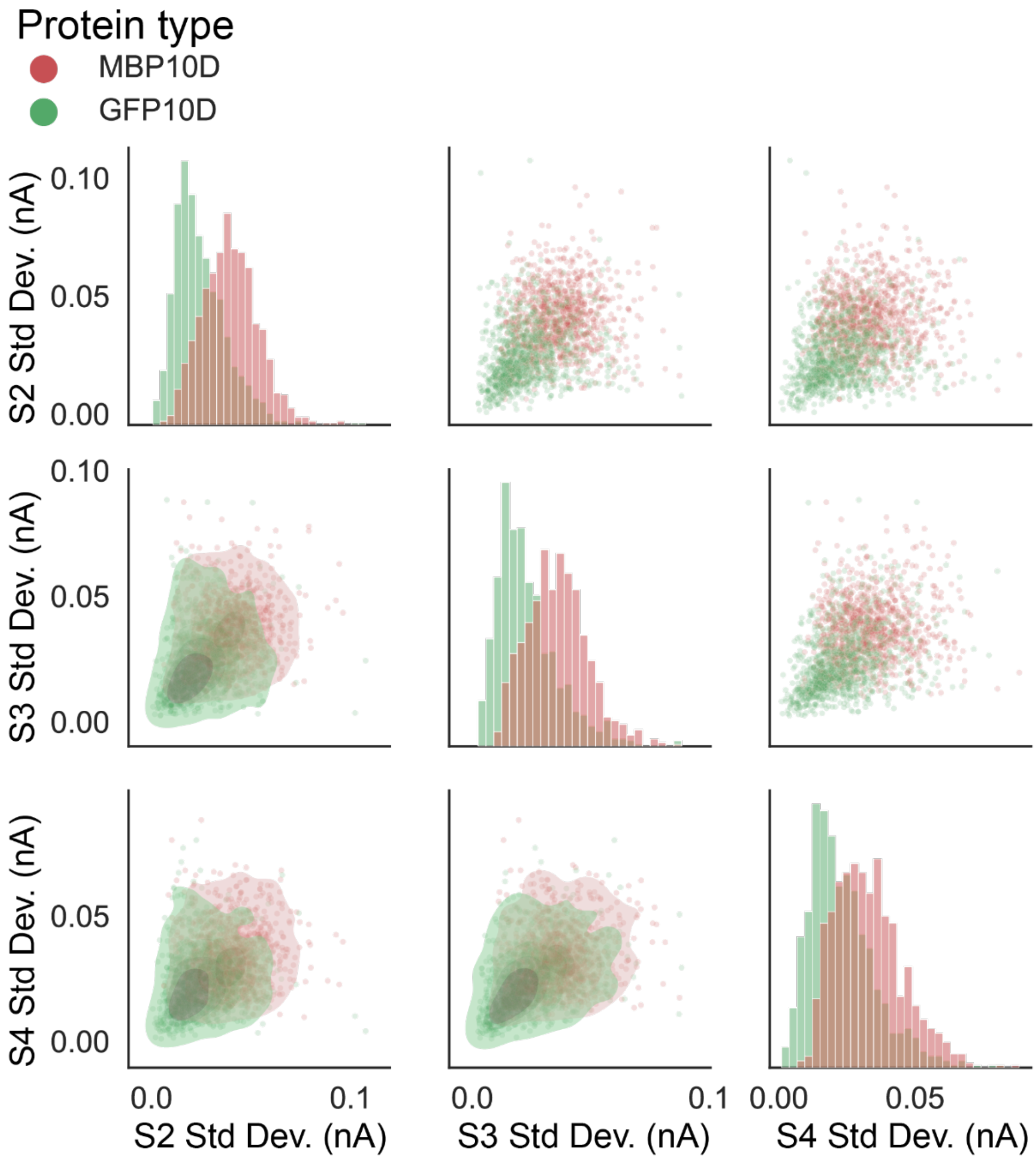


**Figure S30.** Separation of top 3 weighted features, current standard deviation of segments S2, S3, and S4, for GBC model fit for discrimination of MBP-D10 (red) and GFP-D10 (green) based on feature importance heatmap shown in **Figure 5F**. Features are compared to each other with 2D scatter plots (upper-half), histogram distribution for each feature (diagonal), and contour plots (lower-half).

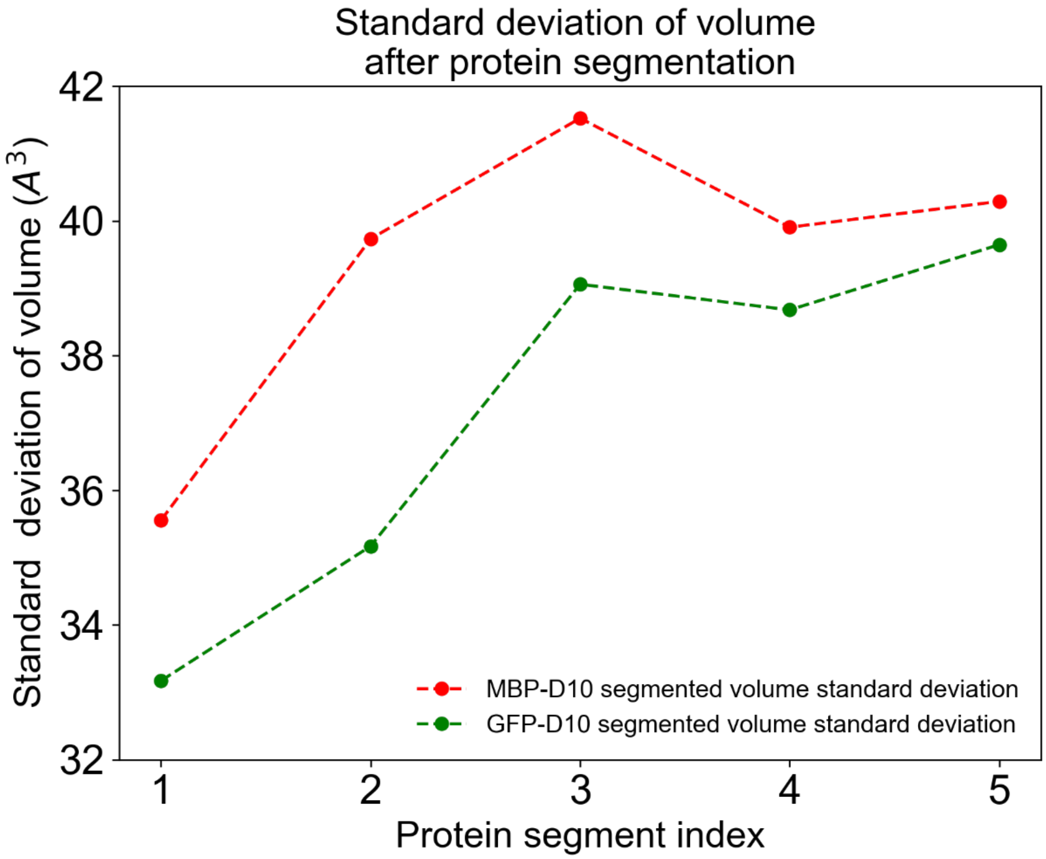


**Figure S31.** Standard deviation of volume after dividing the amino acid sequence for MBP-D10 (389 aa), shown in red, and GFP-D10 (254 aa), shown in green, into 5 segments of equal length.

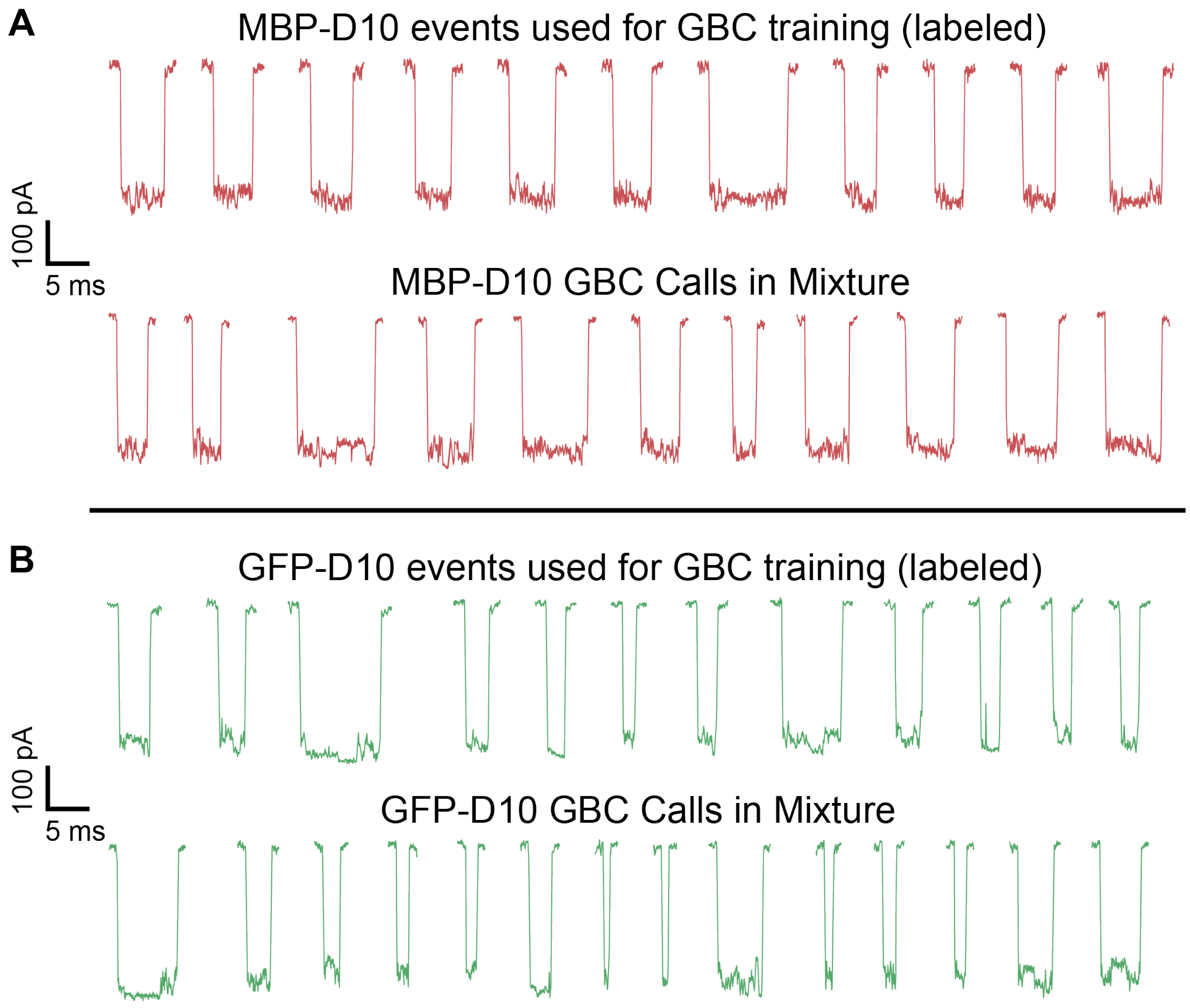


**Figure S32.** Event traces from pure MBP-D10 and GFP-D10 experiments used for training GBC model, where 51 input features from each event were used to construct GBC decision boundaries (upper). Traces of unlabeled translocation events from mixture experiment that were classified by trained GBC model (lower). All experiments were performed in 1.0 M KCl, 2.0 M GdmCl, 10 mM Tris, pH 7.5 with 175 mV bias applied to the *trans* chamber.

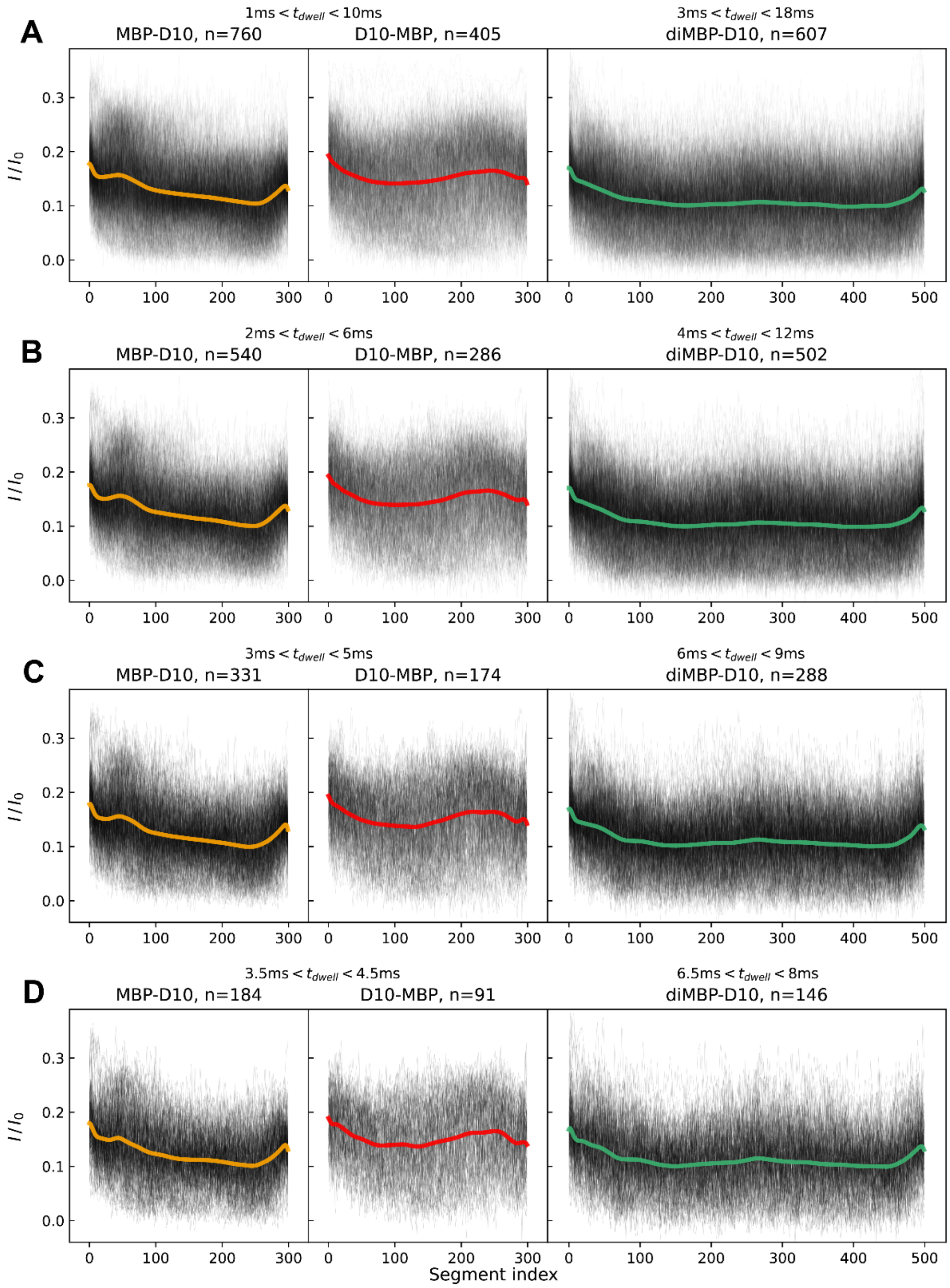


**Figure S33.** Soft-DTW barycenter computation performed on event populations selected via different dwell-time restriction criteria. Monomer variants (MBP-D10 and D10-MBP) share the same selection range in each set. From **A** to **D**, the range goes from wide to narrow. The dwell time range ranges in panel **C** were used in **Figure 4A** of the manuscript.

#### Supplementary Note 3: Soft-DTW Analysis

Turning to the signal content, we first analyzed the three cases where N-terminus vs. C-terminus entry of MBP were studied. The datasets used were MBP-D10, D10-MBP, and diMBP-D10 (all 2 M GdmCl, 1 M KCl, 175 mV). To extract an "average shape" for events of each variant, barycenter computation (Fréchet means) was used. A barycenter or Fréchet mean is a new target time-series $\vec{x}$ that produces the minimum sum of "distance metrics" to all elements $\vec{y}_{1 \dots m}$ in a time-series dataset $Y$ evaluated one-by-one:

$$\min_{\vec{x} \in R^n} \sum_{i=1}^{m} \frac{1}{n \cdot m} \mathbf{dtw}_{\gamma}(\vec{x}, \vec{y_i}) \qquad \textbf{(Eq. S3)}$$

where each $\vec{y}_i$ is an event's current array divided by the open pore current value $I_0$ to normalize for slight variations among different experiments. $n$ is the number of points in each time series, in this case resampled to a fixed length of 300 for MBP-D10 and D10-MBP, and 500 for diMBP-D10. The function $\mathbf{dtw}_{\gamma}$ provides a smoothed (soft) dynamic time warping distance value (loss) computed for the best alignment between the two time-series inputs with the smoothing parameter $\gamma$[6]. We also looked into barycenter computation with the classic dynamic time warping loss function which uses an alignment algorithm based on simple Euclidean distance minimization, but the resulting curve was noisy and could not inform our analysis.

The result of our soft-DTW barycenter computation is a smooth curve, representing the centroid or the "essence" of the translocation events in the dataset. The analysis was performed using tools provided in the tslearn library (v 0.5.2)[7]. This computation was repeated for the three protein variants mentioned above. **Table S4** shows the restrictions applied to select the events, number of passing events, and the suggested smoothing parameter $\gamma$ calculated for each variant set using tslearn's tools. The dwell-time-based selection left out the tails of the variant distributions, limiting the degree of variability in translocation velocity among each variant's population. This was especially important to identify "W" shape and the local maxima (bumps) of the barycenter of diMBP-D10, which would otherwise appear somewhat flat under a wider dwell-time range (refer to **Figure S33** for comparison of various dwell time ranges and their respective barycenters).

The barycenters $\vec{x}_{MBP-D10}$, $\vec{x}_{D10-MBP}$, $\vec{x}_{diMBP-D10}$ are shown in **Figure 4A** along with their respective resampled events in the background. These curves show a smooth trend of how the events of each protein variant tend to progress on average. While both monomer samples show an overall downward trend in their barycenters, they distinctly have opposite locations for local maxima and minima. This DTW method was used to demonstrate unidirectional translocation. For further validation of unidirectional transport, we expected the barycenter of diMBP-D10 to appear more like the MBP-D10 curve, just repeated twice. That was indeed true, as indicated by the initial bump in the beginning of the diMBP-D10 curve which is similar to MBP-D10 and opposite of D10-MBP. While the same soft-DTW metric function can be used for classification purposes, it was omitted in this study to keep the classification features used in the GBC classification analysis simple, understandable, and computationally efficient.

We note that a similar "average shape" analysis could have been performed using signal segmentation and segment averaging across all events of each protein type. However, the time-domain variations across tens or hundreds of events would lead to a highly flat and less informative average shape. This is mainly due to variations in translocation kinetics. The DTW analysis allowed us to minimize the effect of random velocity variations and focus on the overall trend when producing the average curve (barycenter).

#### Supplementary Note 4: Gradient Boosting Classifiers Analysis

##### Overview

To explore whether the signal properties are reproducible and comprise sufficient information to distinguish the translocation events of different proteins, two gradient boosting classifiers (GBC) were trained to discriminate 1) MBP-D10, D10-MBP, and diMBP-D10, denoted as MBP variants (**Figure 4**) for three-way classification and 2) MBP-D10 and GFP-D10 (**Figure 5**) for two-way classification of different types of proteins. Each GBC model was trained and tested with time-domain signal features extracted from translocation events recorded during experiments with a single type of protein, which we define as our labeled dataset. The GBC classifier was built using the *GradientBoostingClassifier* module provided by the scikit-learn (v 1.0.2) python library[8]. The performance of the GBC model generated for classification of MBP variants was assessed with a test set comprised of labeled samples that were randomly chosen and removed from the dataframe containing the training samples used to fit the parameters of the classifier. In addition to measuring the performance of the GBC model built for MBP-D10 and GFP-D10 discrimination, we applied the trained model to classify translocation events from mixture experiments with varying concentration of MBP-D10 to GFP-D10. We define the mixture experiments as our unlabeled dataset. To mitigate the introduction of experimental noise into the model, all events used for this analysis were recorded with 2 M GdmCl, 1 M KCl, and +175 mV bias. Moreover, all training features were divided by the experiment's open-pore current, $I_0$, prior to normalization.

##### Feature Extraction

Event extraction was done with a filter-derivative parsing method to remove event start and end borders (drops), allowing for meaningful computation of event features used for model training (e.g. local minima and maxima ionic current measurements). Every model generated for ML-based classification was fit with a feature space composed of 70 statistical parameters corresponding to segment information within a translocation event. Specifically, each event used for GBC training and testing was divided into 10 segments of equal length and for each segment the 1) minimum, 2) maximum, 3) mean, 4) standard deviation, 5) first quartile (25%), 6) median, and 7) third quartile (75%) were computed, providing a total of 70 features per event. Dwell time was not considered as an input feature to observe if proteins with or without overlapping durations could be resolved.

##### Data preprocessing

Translocation events with dwell times ranging between 0.3 and 20 ms were considered for ML analysis. Sample size of events that met the dwell time conditions for each protein type can be seen in **Table S5**. Segment-level features of all three protein variants (MBP-D10, D10-MBP, and diMBP-D10) and GFP-D10 events were compiled into a dataframe. Next, an equal number of samples for each protein class was prepared to train and test each GBC model on a balanced dataset, the sample size for each protein class was limited by the one with the smallest sample size. For example, the GBC model generated for discrimination of MBP variants was built with 504 samples of MBP-D10, D10-MBP, and diMBP-D10 since we were limited by the number of D10-MBP events. The 504 samples selected for MBP-D10 and D10-MBP were randomly sampled from the total dataset every time prior to GBC training and testing. After creating a balanced dataset, 80% of the data was binned into the training set to fit the GBC and the remaining 20% was withheld to test the accuracy of the trained model. Prior to building the GBC model, the training set was normalized with scikit-learn's *StandardScaler* function, where the mean was centered around 0 and the first standard deviation was +/-1. The normalization parameters for the training set were then used to scale the data in the test set (20%) and data extracted from the unlabeled MBP-D10 and GFP-D10 mixture experiments.

**Model training/testing/evaluation/feature importance**

Both GBC models were built with the command *GradientBoostingClassifier (n_estimators = 200, learning_rate = 0.1, max_features = 70, max_depth = 5)*. The dataset was subjected to a unique and random train-test split prior to building the model, which was repeated 9 times to get an average performance of the model accuracy. The classification results for each individual model generated for this analysis are shown as confusion matrices in **Figures S28** (three-way GBC classifier for MBP variants), and **Figure S29** (two-way GBC classifier for MBP-D10 and GFP-D10), with the mean and standard deviation of all 9 models shown in **Figure 4B** and **Figure 5E**, respectively. Next, the 9 GBC models generated for discrimination of MBP-D10 and GFP-D10 were used to classify unlabeled events extracted from mixture experiments with varying ratios of MBP-D10 and GFP-D10 that had dwell times ranging from 0.3 ms to 20 ms. Shown in **Table S6** are the sample size of events extracted from each ratio experiment under the dwell time conditions. The ratio of MBP-D10 to GFP-D10 called by the 9 GBC models are shown in **Figure 5D** (mean +/- standard deviation) and **Table S7**. Note, there is difference between 50:50 ratio Exp. 1 and Exp. 2 results, which we believe is due to experimental errors, such as pipetting, and not the trained GBC model. Shown in **Figures 5A-C** are example traces for MBP-D10 to GFP-D10 for 20:80, 50:50, and 80:20 ratio experiments with the associated probability scores for each event classified by a trained GBC model. The probability score of each called event in the mixture experiment was determined with sklearn's *predict_proba* module. The mean and standard deviation probability scores determined by the model for all MBP-D10 and GFP-D10 classified events for each mixture experiment is shown in **Table S7**. Finally, to determine which features, i.e., portions of the signal, contribute to the accuracy of the GBC model, we used scikit-learn's *feature_importance* tool to obtain the weights for all 70 features used to fit both GBC models. Simply, the function provides an estimate of the relative importance for each feature during model fitting. The heatmap shown in **Figure 4C** (GBC model built for classification of MBP variants) and **Figure 5F** (GBC model built for classification of MBP-D10 and GFP-D10) show the relative feature importance for each model parameter (normalized). The values (in total add up to 100) and "heat" observed in each box represents the average weighted importance of all 9 GBC models trained and tested for this analysis. Based on **Figure S30**, MBP-D10 shows a higher standard deviation at these segments, providing sufficient separation between the two proteins. Interestingly, if we bin the amino acid sequence of both proteins into 5 equal segments (shown in **Figure S31**), we observe MBP-D10 and GFP-D10 to have the greatest difference in standard deviation at segment 2, which is likely to correspond to event segments 2, 3, and 4. While a segmentation size of 10 was an arbitrary decision, it allowed us to confirm that our system can reproducibly capture distinct trends generated by the translocation of specific sequence contexts through the pore constriction.
